# Prey age modifies risk-taking behavior in a multi-predator environment

**DOI:** 10.64898/2026.07.21.739907

**Authors:** Brian J. Smith, Tal Avgar, Scott D. Peacor, Daniel R. Stahler, Matthew C. Metz, Jack W. Rabe, Wesley Binder, Daniel R. MacNulty

## Abstract

Many species of animals undergo senescence, impacting predator-prey relationships, yet how senescing prey adjust their risk-taking behavior is poorly understood. We used integrated step selection analysis to quantify risk-taking from empirical data. A graphical framework of age-dependent adaptive risk-taking predicted – and our empirical analysis found – a reduction in female elk (*Cervus canadensis*) risk-taking with age toward wolves (*Canis lupus*) but not cougars (*Puma concolor*), underscoring the role of predator hunting mode in ecological dynamics. We estimated average risk-taking toward wolves would be 42% lower in a population with median age 10 versus 4 years, highlighting how a prey population’s age structure likely impacts risk-induced trait responses and the potential emergence of predation-risk effects. Our findings suggest that altered risk-taking may be an adaptive behavioral shift rather than a passive consequence of physical decline. Such adaptive changes to risk-taking may represent an underappreciated link between individual behavior and community-level coexistence.

## Introduction

Senescence is widespread in wild animal populations (Nussey *et al*. 2013) and often manifests as reduced survival (Froy *et al*. 2018), reproduction (Lemaître & Gaillard 2017), and physical ability (MacNulty *et al*. 2009). Although age is known to influence population-level processes (Hoy *et al*. 2019; Jackson *et al*. 2020), its broader ecological consequences, including its effects on species interactions, remain far less understood (Miller & Rudolf 2011). Classic studies of predator-prey interactions show that senescence increases prey vulnerability to predation with implications for predator-prey coexistence (Murdoch *et al*. 2003; Slobodkin 1968). However, more recent research on predation-risk effects, including the risk-induced changes in plastic traits that prey adopt to avoid being eaten (Peacor *et al*. 2020), has largely ignored the role of aging. This gap is important because unrecognized age-dependent variation in risk effects may contribute to uncertainty about the general relevance of risk effects in natural, unmanipulated communities (Peacor *et al*. 2022). Indeed, debates about the ecological significance of risk effects center on long-lived animals that undergo biological aging such as mammalian carnivores and ungulates (Moll *et al*. 2016; Prugh *et al*. 2019).

One line of research addressing how age influences predation risk effects has focused on ontogenesis. For example, many studies of amphibians and insects have examined how vulnerable larvae metamorphose into relatively invulnerable adults, and how plasticity in the timing of this transition can shape demographic outcomes (reviewed in Benard 2004). However, in many species, relatively invulnerable adults undergo a second transition to renewed vulnerability during senescence. Understanding how senescing individuals alter their traits in response to predation risk is an important, yet largely unexplored, aspect of community ecology.

We investigated how senescence affects risk-taking behavior (foraging despite risk) by adult female elk (*Cervus canadensis*) in northern Yellowstone National Park (YNP), USA. Elk have a slow life-history strategy and are known to face tradeoffs between survival and reproduction when forage is limiting (Morano *et al*. 2013). Like other large herbivores, they exhibit high adult survival, with variable reproductive success largely driving their population dynamics (Gaillard *et al*. 1998). Elk in YNP and elsewhere are known to undergo both actuarial and reproductive senescence (MacNulty *et al*. 2020; Smith 2021). During winter, elk inside YNP face predation risk from two predators with different hunting strategies: coursing wolves (*Canis lupus*) and stalking cougars (*Puma concolor*) (Smith *et al*. 2023). Age-related changes in predation risk differ between these predators, with wolves increasing kills of senescent elk more than cougars in YNP (Ruth *et al*. 2019) and elsewhere (Horne *et al*. 2019).

We used a graphical framework (Peacor *et al*. 2013) to capture our assumptions about these changes and develop predictions of how elk should change their defensive investment with age (Box 1). Following Peacor et al. (2013), we expected prey to adjust the expression of a plastic defensive trait to maximize the difference between the resulting benefits (reduced likelihood of being killed by a predator) and costs – reduced “growth” (Fig. 1). By “growth” we refer to all aspects of fitness other than mortality from the focal predator, including mortality from other causes, somatic growth, and reproduction. As defensive trait expression increases, mortality from the focal predator declines, but so does growth (due to reduced energy intake, increased energy expenditure, or increased exposure to other mortality factors). The optimal (hence expected) value of the defensive trait is the maximum of the fitness curve (yellow circles in Fig. 1); the point at which the marginal cost of increased defense exceeds its marginal benefit.

We built on this conceptual framing by further hypothesizing that as prey age and become more vulnerable, the likelihood that they will succumb to a predator’s attack increases, and thus the relative benefits of expressing the defensive trait also increase. However, the degree to which prey vulnerability increases with age depends on the predator and its hunting mode. If predator lethality is mostly independent of prey agility or frailty (thought to be the case for cougars), there should be little change in prey vulnerability with age. If, on the other hand, the predator relies on active pursuit of its prey (as wolves do), aging prey should be significantly more vulnerable. To summarize, age-dependent and predator-specific shifts in the benefits and/or costs of defensive trait expression are expected to result in age-dependent and predator-specific shifts in optimal defensive trait expression.

Our framework makes several predictions. In absolute terms, cougars are likely more lethal than wolves given an encounter, and ambush or stalk-and-chase predators like cougars may induce stronger trait responses than coursing predators like wolves (Schmitz 2005; Schmitz *et al*. 2004). Both of these mechanisms suggest that cougar predation risk should elicit a stronger average lifetime response than wolf predation risk, consistent with prior findings in this system (Kohl *et al*. 2019; Smith *et al*. 2023). However, in terms of relative changes in predator-specific risk-taking with age, we expect that fitness-maximizing aging elk should decrease risk-taking with wolves, but risk-taking with cougars should either remain flat or slightly increase (see Fig. 1E).

We tested these predictions about elk risk-taking behavior in YNP using GPS movement data for elk, wolves, and cougars. To estimate risk-taking behavior, we quantified adult female elk preference for high-risk, high-reward habitat (HRH) versus low-risk, low-reward habitat (LRH) using an integrated step selection analysis (iSSA; Avgar *et al*. 2016). We modeled habitat preference as a function of elk age, while accounting for potentially confounding variables. Relative to younger elk, we found that older elk exhibited less risk-taking with wolves, but not with cougars. Finally, we used these models of risk-taking to illustrate population-level patterns of risk-taking under different age structures, highlighting the broader ecological importance of age-related behavioral change. Our results demonstrate the importance of age for shaping risk effects in long-lived prey species.

## Methods

### Analysis overview

Our analysis consisted of a two-stage modeling procedure, followed by model-based predictions, with uncertainty propagated using parametric bootstrapping (Fig. S5.1). To estimate elk risk-taking behavior, we considered hourly GPS data from 72 adult female elk in YNP over five winters from 2016 – 2020 (hereafter, years). Elk ranged in age from 1.5 – 20.5 y, and 41 individuals were tracked for multiple years (max = 4 y). We only used data corresponding to intensive periods of predation monitoring (hereafter, seasons): early winter (November 15 – December 15) and late winter (March 1 – March 31). We refer to each winter using the year of that January, e.g., 2016 refers to November 15, 2015 – March 31, 2016. The dataset consisted of n = 214 elk-year-seasons. At least 10 elk-year-seasons were available for all ages between 3.5 y and 17.5 y (Fig. S5.2).

The first stage of modeling was to parameterize movement models for each elk-year-season using integrated step-selection analysis (Avgar *et al*. 2016). The fitted integrated step-selection functions (iSSFs) provided our estimate of elk risk-taking behavior by predicting their response to the tradeoff between foraging and predator-specific predation risk (see below). For each elk-year-season iSSF, we predicted the preference for HRH over LRH for each predator. This metric was our estimate of the elk’s risk-taking behavior during that year-season; i.e., it is the elk’s expression of the defensive trait (Box 1).

We used a second stage of modeling to estimate the effect of covariates on the defensive trait at the population level. We modeled defensive trait expression as a function of the elk’s age, while controlling for moderator variables, including elk density, predator density, individual predation risk exposure, season, and year. We used these fitted models to predict population-average trait values for a given age structure, corresponding to the estimated age structure of this population in 1995 (median female age = 4 y) and in 2009 (median female age = 10 y).

### Estimating predation risk

We estimated our key metric, predation risk, by developing a dynamic landscape of risk (dLOR), in which risk varies in space and time (Kohl *et al*. 2018; Palmer *et al*. 2022). We modeled risky places based on locations of elk kills by either wolves or cougars as a function of landscape and temporal covariates (Appendix S1). The resulting risk layers were conditioned on constant elk and predator density; thus, the units were kills/elk/predator (see Appendix S1). We modeled wolves (Fig. 2 A–B) and cougars (Fig. 2 C–D) separately, and we modeled early winter and late winter separately (rows in Fig. 2). We modeled risky times based on the movement rates of GPS-collared wolves and cougars, which we used to develop activity indices for both predators in early (Fig. 2E) and late (Fig. 2F) winter (Appendix S2). The elk iSSA captured risk as an interaction between risky places and risky times, thus creating the dLOR.

**Fig. 2.**
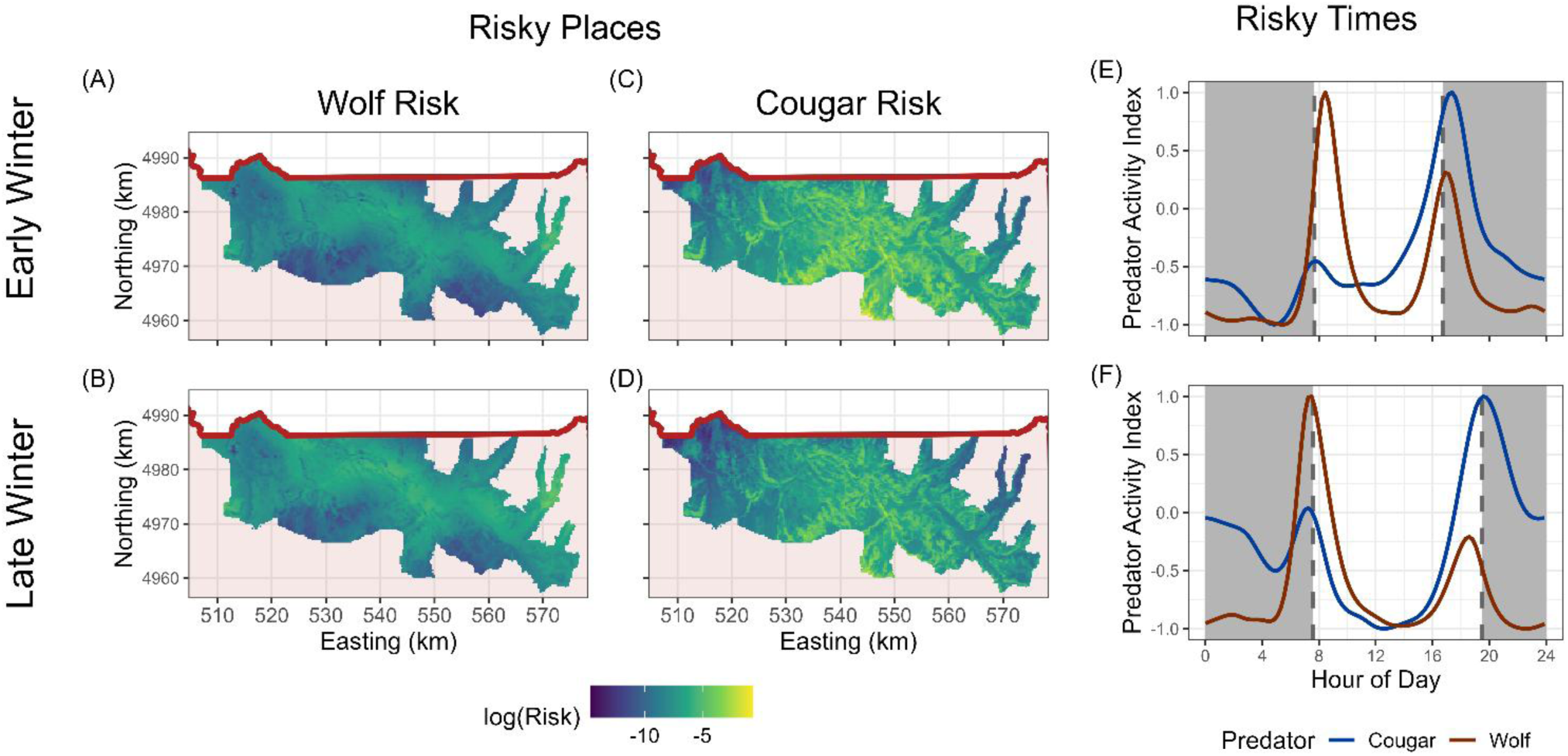
Dynamic landscape of risk. We developed a dynamic landscape of risk by combining separate estimates of risk in space [(**A**) – (**D**)] and time [(**E**) – (**F**)]. We estimated risky places using locations of predator-killed elk in northern Yellowstone. We made predictions for each year from 2016 – 2020; 2020 is shown here, but all years were similar. Units are expected elk kills/elk/predator (shown log-transformed). We estimated risky times from movement rates of collared predators and scaled predicted movement rates from -1 to +1. In [(**A**) – (**D**)], shaded red polygon shows Yellowstone National Park and the risk raster is displayed for the portion of the northern range inside the park. In [(**E**) – (**F**)], shaded areas show nighttime and unshaded areas show daytime. Sunrise and sunset, shown as vertical dashed lines, are for December 1 (**E**) and March 15 (**F**).

### Integrated step-selection analysis

We fit an iSSF for each elk-year-season of data with a suite of covariates (Table 1, Appendix S3). Briefly, we modeled habitat selection as a function of resources (forage biomass), risks (the dLOR), and conditions (canopy openness and terrain roughness) (Matthiopoulos *et al*. 2015). We modeled elk movement as a function of snow-water equivalent and time of day, and we allowed for correlation between step lengths and turn angles. We estimated forage biomass using the herbaceous biomass product from the Rangeland Analysis Platform (Jones *et al*. 2021; Robinson *et al*. 2019; Smith *et al*. 2023), and we allowed selection for biomass to vary with solar time, reflecting the diel pattern in elk feeding. We included openness and roughness, which have been used to model risky places in this system (Kohl *et al*. 2018; Smith *et al*. 2023), and they are included in our risk layers (Appendix S1). We included these landscape variables in the iSSA because elk may also select them for other reasons; e.g., openness facilitates social interactions (Brennan *et al*. 2015) and roughness may create preferred bedding sites (Bender *et al*. 2012).

**Table 1.** Covariates included in the integrated step selection analysis. Habitat selection variables were either resources, risks, or conditions (Matthiopoulos *et al*. 2015). Movement variables update the step-length (Gamma) and turn-angle (von Mises) distributions (Avgar *et al*. 2016). For full model formula, see Appendix S3.

| Category | Covariate | Description |
| --- | --- | --- |
| Resources | Biomass | Herbaceous (grasses and forbs) biomass (kg/ha) estimated by the Rangeland Analysis Platform. Interacted with sine and cosine transformations of solar time of day to allow for diel pattern in foraging. |
| Risks | Dynamic Landscape of Risk (dLOR) | Estimated risk layer for each predator (wolves, cougars), interacted with the diel activity pattern of that predator. See main text and Appendix S1, S2. |
| Conditions | Canopy Openness | Percent of raster pixel *not* covered by trees estimated by the Rangeland Analysis Platform. |
|  | Terrain Roughness | Largest change in elevation (m) among 8 nearest neighbors of focal raster pixel. |
| Movement | Step Length | Updates the scale parameter of the Gamma distribution describing step lengths. |
|  | log(Step Length) | Updates the shape parameter of the Gamma distribution describing step lengths. Interacts with SWE, solar time, and cos(Turn Angle). |
|  | cos(Turn Angle) | Updates the concentration parameter of the von Mises distribution describing turn angles. Interacts with log(Step Length) to allow correlation between step length and turn angle. |
|  | Snow-Water Equivalent (SWE) | SWE (kg/m<sup>2</sup>) from Daymet v4.0. Included in interaction with log(Step Length) to allow step length distribution to change with SWE. |
| Other | cos(Solar Time),<br>sin(Solar Time) | Circular transformations of time of day, based on solar position given the latitude, longitude, and date. Interactions with these terms allows diel pattern in foraging and movement. See Appendix S3. |

### iSSA model validation

We evaluated calibration, discrimination, and explanatory power for each iSSF. We evaluated calibration visually using used-habitat calibration (UHC) plots (Fieberg *et al*. 2018). We measured discrimination using concordance (Harrell *et al*. 1982), estimating the model’s ability to distinguish between used and available steps. We evaluated explanatory power using the measure of explained randomness (MER; O’Quigley *et al*. 2005) – a pseudo-R^2^ considering only the used steps (which are data) and not the available steps (which are for estimation; see Michelot *et al*. 2024).

### Estimating the defensive trait

We took a novel approach to estimating risk-taking behavior using iSSFs. We quantified the defensive trait using relative selection strengths (RSS; Avgar *et al*. 2017), and we presented most results as the natural logarithm of RSS (log-RSS). A value of log-RSS > 0 indicates preference for HRH over LRH, log-RSS < 0 indicates preference for LRH over HRH, and log-RSS = 0 means no preference.

We defined HRH as a habitat with the 90th quantile of biomass and the 90th quantile of risk and LRH as a habitat with the 10th quantile of biomass and the 10th quantile of risk. Only the magnitude, not the sign or significance, of the effect was sensitive to the quantiles of forage and risk we chose. We explain this in detail in Appendix S3, but briefly, the log-RSS for a habitat variable is the fitted coefficient times the difference in the habitat variable. The choice of the quantiles affects the difference in the habitat variable, but the sign and significance are determined by the point estimate and uncertainty in the estimated coefficients.

Because we modeled selection for resources and risk as temporally dynamic (Fig. 3A–F), we made this prediction for each of the 24 h of the day in each season. We summarized this daily variation in risk-taking into three quantities: riskiest time (log-RSS for the hour the predator was most active), safest time (log-RSS for the hour the predator was least active), and daily mean (mean log-RSS across all hours). These three quantities captured the two extremes of risk (riskiest time and safest time) as well as the mean across the day to try to provide a complete view of the variation in risk-taking.

**Fig. 3.**
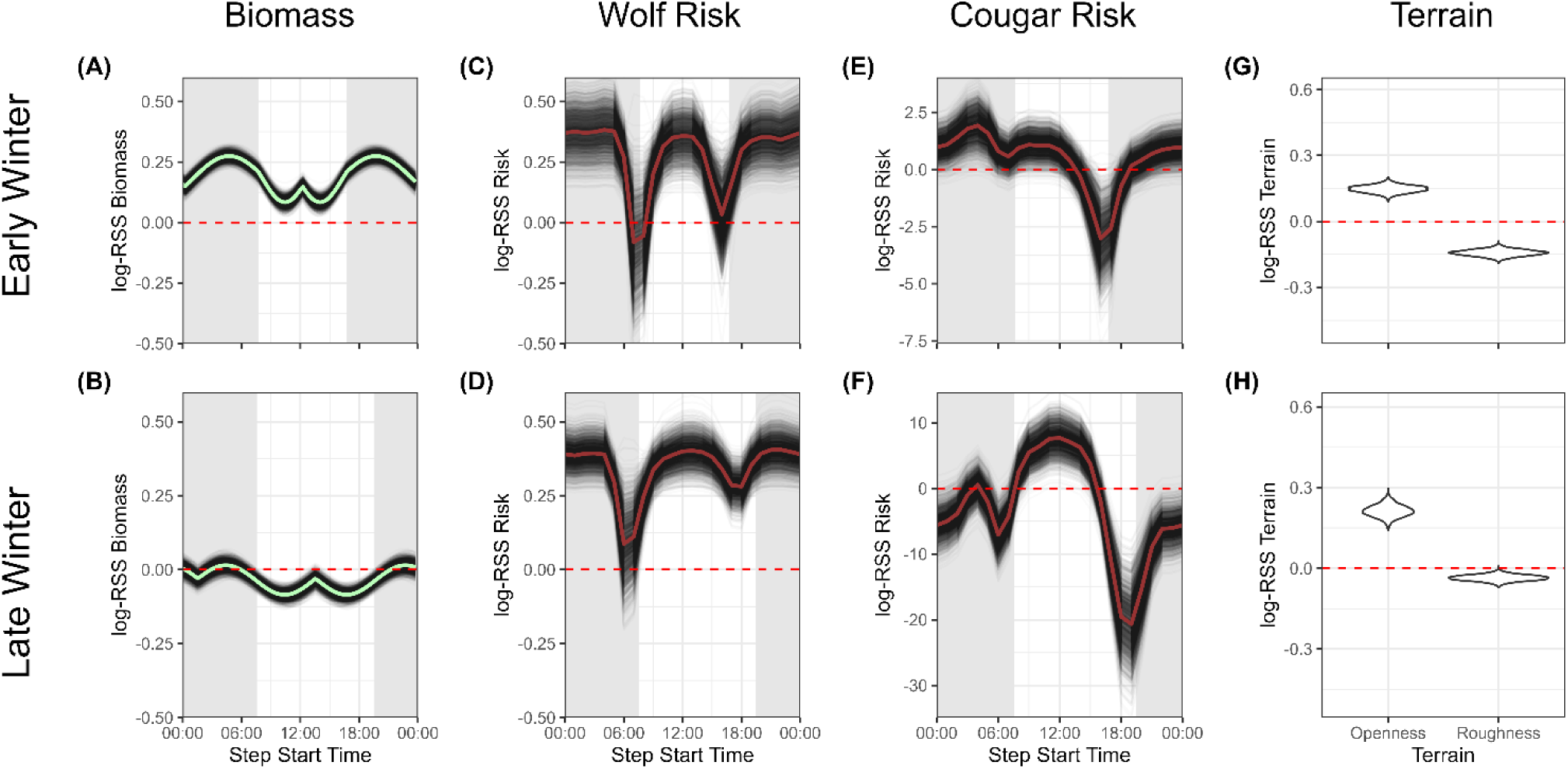
Mean population-level habitat selection. Results from the iSSA, summarized across individuals. Elk selected for biomass at all times of day in early winter (**A**) but showed nearly no selection for biomass during late winter (**B**), possibly indicating forage depletion or snow accumulation limiting access to forage. Elk typically selected for places with wolf risk [(**C**) – (**D**)], except when wolves were most active (see Fig. 2) in early winter (**C**); however, elk typically strongly avoided places with high cougar risk when cougars were active [see Fig. 2; (**E**) – (**F**)]. Elk selected places with more open canopy and less rough terrain in both seasons (**G**) – (**H**). Note that the y-axes in panels (**E**) – (**F**) differ from all other panels. Thick lines in panels (**A**) – (**F**) show the mean across all bootstrap iterations (n = 2000), while the thin, transparent lines show bootstrap iterations. Panels (**G**) and (**H**) show the density across bootstrap iterations. In (**A**) – (**F**), shaded regions show nighttime and unshaded regions show daytime.

### Age-dependent risk-taking

We used generalized additive mixed models (GAMMs) to estimate the relationship between age and these metrics of risk-taking, while controlling for potential moderator variables (see Avgar *et al*. 2020; Winter *et al*. 2024 for details about context dependency in habitat selection). In addition to age, we modeled risk-taking as a function of elk abundance (density dependence), focal predator density (risk intensity), total risk exposure (availability dependence), and season (early or late winter), with random effects for year and individual elk to account for repeated measures. We propagated the uncertainty from the first stage (iSSFs) to the second stage (GAMMs) using parametric bootstrapping, with 2000 iterations. For each bootstrap iteration, we resampled the coefficients of each iSSF from a multivariate normal distribution with mean given by the maximum likelihood coefficients and variance-covariance matrix estimated by the model. We used those new coefficients to estimate the behavioral trait (log-RSS) for each iteration, fit the GAMMs, and calculate any derived quantities.

### Population-level outcomes

We used the bootstrapped GAMMs to predict, for a given age structure, the population-mean risk-taking behavior. We altered only the age structure, holding all other covariates at their mean. We used the age-class reconstruction of Hoy et al. (2019) to choose age structures this population had experienced in recent decades. We chose years representing the youngest median age (4 y, 1995) and the oldest median age (10 y, 2009) in the reconstruction of Hoy et al. (2019). We predicted the trait value for each age from 1 – 20, then took the mean, weighted by the proportion of the population in that age class under the reconstruction.

## Results

### Integrated step-selection analysis

The spatial correlation between forage biomass and wolf risk was weakly positive (Spearman’s r = 0.17; Fig. S5.3A), while the spatial correlation between forage biomass and cougar risk was weakly negative (Spearman’s r = -0.16; Fig. S5.3B). Elk selection for forage biomass shifted with (solar) time of day (Fig. 3A). Elk generally selected areas with higher biomass (accumulated over the preceding growing season) during early winter (Fig. 3A), but this selection weakened in late winter (Fig. 3B), possibly reflecting seasonal depletion of forage or snow accumulation limiting forage access. Elk generally avoided high-risk areas when predators were most active (troughs in Fig. 3C–F) and selected them during predator downtimes (peaks in Fig. 3C–F), consistent with temporal risk allocation strategies that allow elk to use high-reward habitat while minimizing predation risk. Elk also selected for open canopy and avoided rough terrain (above and beyond the effects of the predation-risk layers; Fig. 3G–H), consistent with broader population-level patterns (Smith *et al*. 2023). As expected, elk step length distributions were largest for turn angles near 0 (Fig. S5.4A–B), decreased with SWE (Fig. S5.4C–D), and varied with time of day such that the longest steps occurred in morning and evening (Fig. S5.4E–F).

### iSSF validation

Visual inspection of UHC plots indicated very good calibration between the models and the observed used habitat (Fig. S5.5). The models had good discrimination between used and available steps, with mean concordance = 0.783 (2.5th quantile = 0.708, 97.5th quantile = 0.861. The models also had moderately strong explanatory power, with MER mean = 0.617 (2.5th quantile = 0.339, 97.5th quantile = 0.821).

### Age-dependent risk-taking

Consistent with our predictions, risk-taking at the riskiest time of day declined with age in response to wolves (Fig. 4A) but increased with age in response to cougars (Fig. 4B). The decline in wolf-related risk-taking was linear, with the youngest elk taking the most risk (Fig. 4A). In contrast, cougar-related risk-taking showed a non-monotonic pattern with prime-aged elk taking the least amount of risk (bootstrap mean minimum = 7.9 y, log-RSS mean = -8.1, SE = 4.5; Fig. 4B). Across all ages, wolf-related risk-taking was near 0 (mean = 0.1, SE = 0.7), indicating that elk were, on average, equally likely to select HRH or LRH (Fig. 4A). In contrast, the average value of cougar-related risk-taking was strongly negative (mean = -6.5, SE = 4.2), indicating elk generally avoided HRH in the presence of cougar risk (Fig. 4B).

**Fig. 4.**
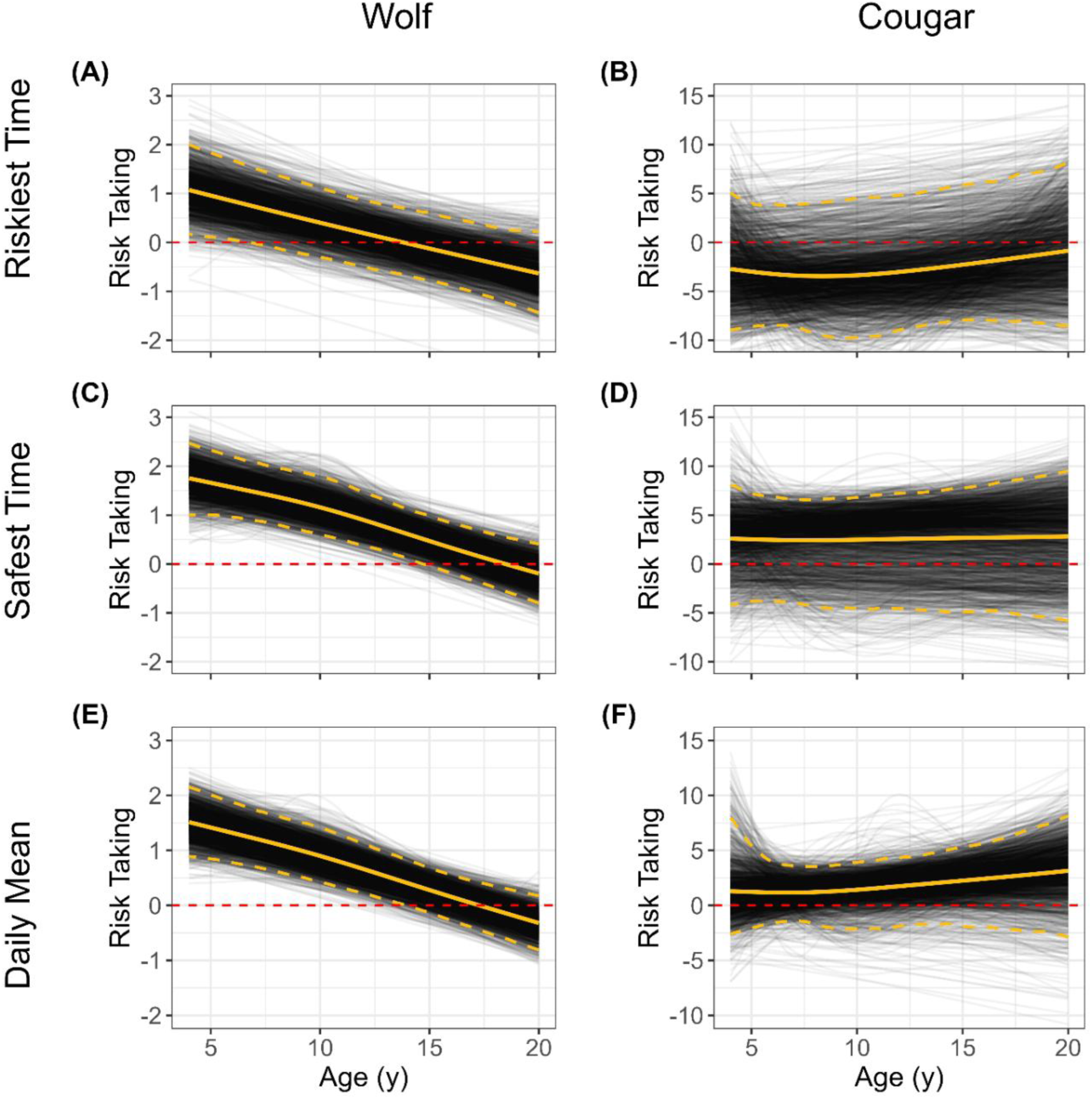
Age-dependent risk-taking. We quantified risk-taking with predictions from each elk-year-season movement model. We defined risk-taking as the log relative selection strength (log-RSS) for a high-risk/high-reward habitat (HRH; 90th quantile biomass, 90th quantile predation risk from focal predator) versus a low-risk/low-reward habitat (LRH; 10th quantile biomass, 10th quantile predation risk). Positive values indicate selection for the HRH whereas negative values indicate selection for the LRH. Selection for biomass and risk changed with time of day, and here we summarize risk-taking at the riskiest time (highest activity of the focal predator), at the safest time (lowest activity of the focal predator), and the daily mean. Risk-taking with respect to wolves was generally positive, but declined with age (**A**, **C**, **E**). Risk-taking with respect to cougars was generally negative at the riskiest time (**B**), with a slight increase with age. It was generally positive at the safest time, with no age pattern (**D**). On average throughout the day, elk selection tended to be slightly positive and not age-related (**F**). Note different y-axis ranges for the wolf and cougar columns. Compare these results to our predictions in Fig. 1E.

Risk-taking during the safest time of day also declined with age in response to wolves (Fig. 4C), but showed little age-related pattern in response to cougars, for which the 95% confidence interval consistently overlapped 0 (Fig. 4D). Averaged across ages, elk preferred the HRH over the LRH during safe times with respect to wolves (mean = 1.3, SE = 0.7; Fig. 4C). With respect to cougars, the mean was also positive, although uncertainty was much greater (mean = 2.3, SE = 2.7; Fig. 4D). The difference in risk-taking between the riskiest and safest times of the day – i.e., between Fig. 4C and 4A for wolves, and Fig. 4D and 4B for cougars – was positive for both predators, indicating that elk were more likely to use risky places during safe times (Kohl *et al*. 2018, 2019). However, this temporal shift in habitat use was more pronounced with respect to cougars (mean difference = 7.50, SE = 4.0) than wolves (mean difference = 1.17, SE = 0.56).

Daily mean risk-taking also declined with age in response to wolves (Fig. 4E) but showed no consistent age-related pattern in response to cougars (Fig. 4F). For adult elk younger than 15 years, daily mean risk-taking with respect to wolves was positive (Fig. 4E), indicating a preference for HRH over LRH. Only after age 15 did this preference diminish to a toss-up (log-RSS = 0). In contrast, elk showed no clear age-based pattern in response to cougars, and the daily average log-RSS remained near zero (Fig. 4F).

### Moderator Variables

Risk-taking with both predators increased with elk density (Fig. S5.6), consistent with positive density-dependent habitat selection (Avgar *et al*. 2020; Smith *et al*. 2023), and more specifically, a “safety in numbers” effect (Lehtonen & Jaatinen 2016). We also found that cumulative risk exposure influenced risk-taking behavior (Fig. S5.7), consistent with availability-dependent habitat selection behavior (“functional response”; Avgar *et al*. 2020; Winter *et al*. 2024). Specifically, risk-taking during the safest time of day decreased with increasing exposure to both wolf and cougar risk (Fig. S5.7C–D); i.e., elk exposed to higher predation risk behaved more cautiously, even at the safest time of day. Averaged across the day, increased exposure to wolf risk led to reduced risk-taking (Fig. S5.7E), whereas exposure to cougar risk had little effect on daily mean behavior (Fig. S5.7F). We found no significant effects of predator density (Fig. S5.8), season (Fig. S5.9), or year (Fig. S5.10) on risk-taking.

### Population-level outcomes

We predicted population-level mean risk-taking for both predators and all three temporal summaries (riskiest time, safest time, daily mean; Table S4.1). However, due to the high uncertainty in age-related risk-taking responses to cougars, we report results here only for wolves. The predicted mean daily risk-taking (log-RSS) for a young elk population (median age = 4 y) was 0.94 (95% CI = 0.81 – 1.97), while for an older population (median age = 10 y; Fig. 5A) it was 0.38 (95% CI = 0.38 – 1.27; Fig. 5B). On the natural scale (RSS), this corresponds to a 42% decrease (95% CI = 28 – 52%) in probability of selecting HRH over LRH. Similar reductions were observed for risk-taking at the riskiest (41%, 95% CI = 23 – 58%) and safest times of day (42%, 95% CI = 25 – 55%).

**Fig. 5.**
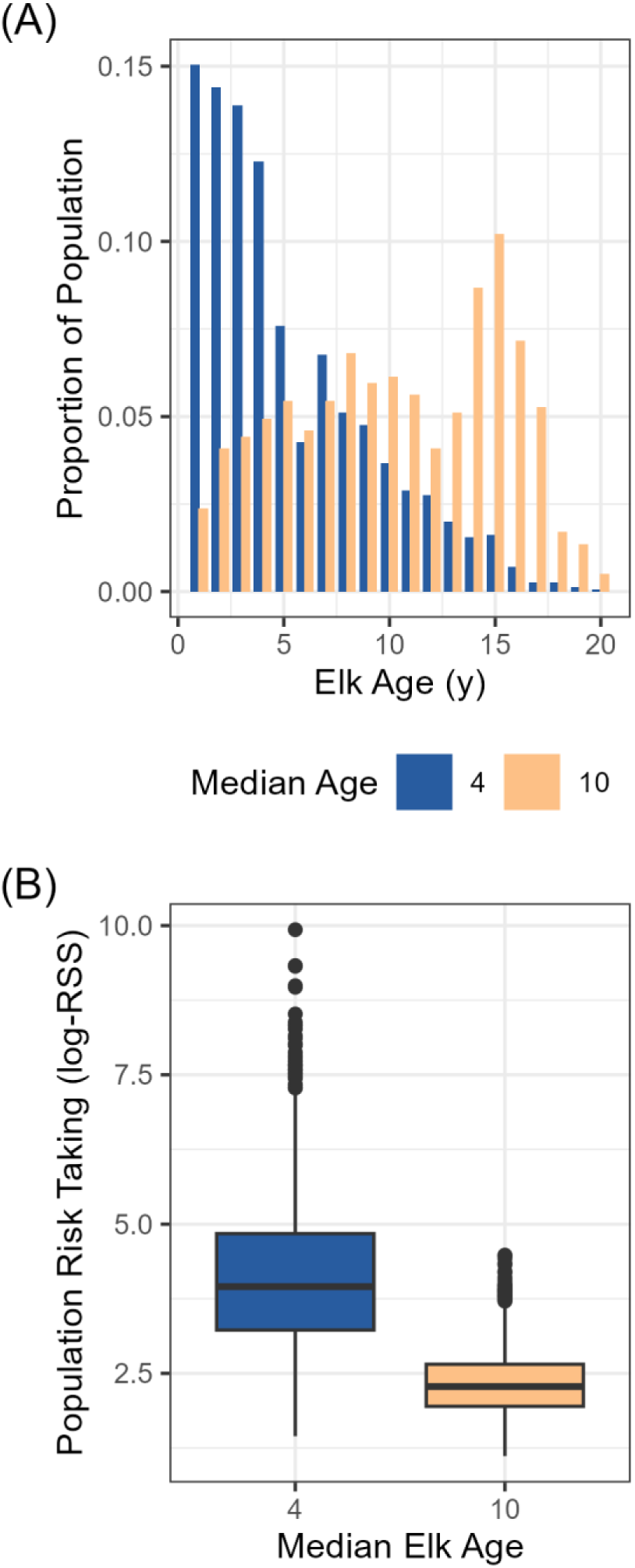
Population-mean risk-taking behavior. We used our fitted model to predict the risk-taking behavior for all age classes of elk in the population (1 – 20). We used the reconstruction of Hoy et al. (2019) to estimate the proportion of each age class in the population for two age structures **(A)**: a relatively young population, with median female age = 4 y (estimated for 1995); and a relatively old population, with median female age = 10 y (estimated for 2009). We used the proportion to take a weighted mean of the risk-taking trait, producing a population-level average specific to an age structure **(B)**. The boxplots show the bootstrap uncertainty around the mean. On the natural scale (RSS), the predicted risk-taking behavior decreased 42% (95% CI = 28 – 52%) as the population aged, with all other predictors held at their mean.

## Discussion

Our results demonstrate that aging plays a central role in shaping risk-induced trait responses in prey. By combining a theoretical framework (Box 1) with detailed statistical analysis, we show that individual age, predator identity, and diel patterns of activity interact to determine how prey forage under risk. This work advances our understanding of the ecological importance of aging beyond population dynamics and direct predation, extending our understanding of how aging affects a key interspecific interaction — a predation-risk effect. Whereas other studies of aging and risk have focused on ontogeny through maturation and adulthood, our work focuses on senescence. Our finding that risk-taking behavior changes with age has implications for understanding when risk effects might emerge in a system, and it elucidates how a prey population’s age structure likely impacts risk-induced trait responses and the potential emergence of nonconsumptive (population) or trait-mediated indirect (community) effects.

Elk exhibited divergent age-related responses to risk from wolves and cougars, consistent with our theoretical predictions (Box 1) and similar to the competing foundational predictions of life history theory. Risk-taking with wolves declined with age, consistent with the reproductive restraint hypothesis (Curio 1983; McNamara *et al*. 2009). In contrast, risk-taking with cougars was flat or slightly increased with age, the latter consistent with the terminal investment hypothesis (Clutton-Brock 1984; Williams 1966). However, these two life-history theory hypotheses (reproductive restraint vs. terminal investment) were developed with a different mechanism from our predictions. The life-history theory approach has largely been framed as a time horizon problem, with organisms at each age navigating a tradeoff between current investment and residual reproductive value (Williams 1966). Whether the time horizon shrinks due to a maximum lifespan or due to somatic damage accumulation, the optimal strategy may be terminal investment or reproductive restraint (McNamara *et al*. 2009). Indeed, the empirical literature shows support for both outcomes across diverse taxa (reviewed in Duffield *et al*. 2017; Jehan *et al*. 2021). Our graphical framework does not consider residual reproductive value, yet it shows how senescence can change reproductive effort resulting in either terminal investment or reproductive restraint. Our framework (Box 1) could be used to capture the value of future reproduction by changing the shapes of the growth and predation curves, and we think this is a fruitful direction for future research integrating predation risk and life-history theory.

Our graphical model (Box 1) also predicts the relative magnitude of the consumptive and nonconsumptive effects (CEs and NCEs) of the predators (Peacor *et al*. 2013). For a given level of the defensive trait, the CE is given by the predation rate, while the NCE is given by the difference from the maximum growth rate. Our model predicts that the NCE for prime-aged elk should be low for both predators but that it should increase for senescent elk. For senescent elk, the predicted relative contribution of the NCE is larger for cougars than for wolves because the reduction in growth rate (NCE) is larger and the predation due to mortality (CE) is lower at the predicted higher expression of the defensive trait. An NCE primarily affecting senescent individuals would have a dampened effect on population growth rate compared with an NCE affecting all ages, emphasizing the importance of age in NCEs.

Our results support previous work showing that cougar risk in this system has a stronger effect on elk habitat selection than wolf risk (Kohl *et al*. 2019; Smith *et al*. 2023). On average, elk avoided HRH more strongly in response to cougars than to wolves, even though cougar risk varied less with age. This asymmetry is consistent with two complementary explanations: (1) cougars are more lethal than wolves given an encounter (Hornocker 1970), and/or (2) stalking predators like cougars elicit stronger trait responses than cursorial predators like wolves (Schmitz *et al*. 2004). In addition, forage biomass was weakly negatively correlated with cougar risk but weakly positively correlated with wolf risk (Fig. S5.3). These opposing relationships suggest that the structure of the foraging–risk trade-off differs across predator types, with cougars imposing strong costs even in lower-reward areas. These patterns collectively suggest that predator identity, not just the presence of predation risk, fundamentally determines prey behavior and its downstream ecological effects.

Although the age effect for cougar-related risk-taking was more uncertain and nonmonotonic, elk of prime age avoided risk the most, whereas older individuals appeared more willing to accept risk, possibly to meet foraging needs late in life. This divergence raises the possibility of compensatory behavior: as aging elk reduce foraging in response to wolf risk, they may offset this cost by increasing foraging under cougar risk. Indeed, senescing mule deer (*Odocoileus hemionus*) can compensate for declining fitness by increasing foraging effort (Levine *et al*. 2026). These findings about cougar risk also suggest an important alternative explanation for the general ecological pattern of elevated predation risk observed in older individuals. Senescent prey are often assumed to be more vulnerable because of deteriorating physical condition, but our findings suggest that heightened risk-taking may be an adaptive behavioral shift rather than a passive consequence of physical decline. In our case, that would suggest that older elk may be more vulnerable to cougar predation (Ruth *et al*. 2019) not only because they are inherently weaker, but because they deliberately take more risks—a pattern consistent with terminal investment. These patterns emerge for both actuarial and reproductive senescence in elk (MacNulty *et al*. 2020; Smith 2021), and our results suggest that behavioral risk responses may reflect alternative life-history strategies that arise when these two components of senescence diverge (Bouwhuis *et al*. 2012).

Elk exhibited strong diel shifts in risk-taking, selectively using risky habitats during periods of reduced predator activity (Fig. 3C-F). This temporal risk allocation was evident for both predators but especially pronounced in response to cougars (compare y-axes in Fig. 3C-D with Fig. 3E-F). Daily mean risk-taking responses masked this behavioral flexibility, reinforcing the importance of considering temporal variability in predation-risk dynamics (Lima & Bednekoff 1999). That elk time their habitat use to avoid high-risk periods likely weakens trait-mediated trophic cascades (Brice *et al*. 2025; Kohl *et al*. 2018), since foraging continues even in risky places but at safe times.

Elk selection for high-biomass habitats weakened in late winter (Fig. 3A-B), likely due to either forage depletion or snow accumulation, which may have altered the reward side of the risk-reward trade-off. Elk on average stopped avoiding wolf risk during this time, even at risky times (Fig. 3D). These seasonal changes suggest elk engage in state-dependent risk-taking, where animals in poorer condition may take more risks to meet immediate nutritional needs (Heithaus et al. 2007). Indeed, female elk entering winter with less body fat are known to lose less fat than fatter females – a pattern thought to be related to physiological compensation (Cook *et al*. 2013) but which our results suggest may also be related to state-dependent risk-taking. Conversely, elk avoidance of cougar risk became even stronger in late winter, especially during night, suggesting that responses are predator specific. Perhaps harsher winter conditions make cougar risk even greater in already risky places.

Elk risk-taking behavior was also shaped by exposure to risk and conspecific density. Elk living in overall riskier places took fewer risks, consistent with the ability to perceive the riskiness of their surroundings. Elk took more risks when conspecific density was high, suggesting a “safety in numbers” effect (Lehtonen & Jaatinen 2016), possibly due to dilution of individual predation risk or increased vigilance from group members. These two effects observed in individual behavior agree with broader patterns seen in the population distribution (Smith *et al*. 2023). Together, these patterns underscore the complexity of behavioral decision-making in natural systems and highlight that risk effects are context dependent — modulated not just by predator presence but also by time, space, age, physiological condition, and social environment.

In contrast, predator density itself had no measurable effect on risk-taking behavior. This is surprising, as clearly predator density is a major factor in the level of risk. Predator density in this system did not change much from 2016 – 2020 (wolf density: 2.77 – 4.62 km^-2^; cougar density: 2.05 – 2.33 km^-2^), which likely influenced our ability to detect a pattern (Fig. S5.8), but it may have also influenced the ability of elk to detect a change. We also found no pattern by season or year, suggesting the predictors in our model already captured important temporal variation.

The ecological consequences of age-structured risk-taking behavior are substantial. Using realistic age distributions from the YNP elk population (Hoy *et al*. 2019), we estimated that population-level risk-taking declined by 42% as the population aged from a median age of 4 y (seen in 1995) to a median age of 10 y (seen in 2009). Because large herbivores like elk have slow life histories and adult female survival is the most elastic vital rate (Raithel et al. 2007), reduced risk-taking in older populations may help stabilize population size in the face of predation. This dynamic may resemble a type III functional response, where predation rates decline at low prey densities due to behavioral or demographic shifts that reduce exposure (Rosenzweig & MacArthur 1963). Moreover, if predators disproportionately remove very young and very old individuals—the age classes with the weakest antipredator responses—this could generate negative feedback in predation rates. As predation shifts the age structure toward more prime-aged individuals with stronger antipredator behavior, overall prey vulnerability would decline, thereby reducing predator success. Such negative eco-demographic feedbacks should be expected to stabilize predator–prey dynamics and represent an underappreciated mechanism linking individual behavior to community-level coexistence (Fryxell & Lundberg 1998).

Our findings emphasize the value of integrating life-history theory and predator–prey ecology to understand risk effects in the wild. Although both reproductive restraint and terminal investment have been observed in other systems (Duffield et al. 2017), our study is among the first to show that these opposing age-based strategies can occur simultaneously in a single prey population, depending on the predator encountered. These dynamics likely extend beyond elk and carnivores: long-lived prey facing multiple predators—such as primates, cetaceans, or large birds—may similarly modulate risk-taking behavior in predator-specific, age-dependent ways. More broadly, age variation in plastic traits may contribute to uncertainty about the generality of risk effects in free-living systems (Peacor *et al*. 2022).

By applying integrated step selection analysis to long-term GPS movement data, we offer a new approach for linking aging and predator-sensitive behavioral plasticity in the wild. Yet the fitness consequences of these shifts remain unknown. In species where adult survival is key to lifetime reproductive success, even modest decreases in risk-taking could improve fitness. Testing these links empirically will be essential for understanding when behavioral plasticity contributes to population resilience—and when it does not.

## Statement of authorship

BJS, DRM, and TA conceived and developed the idea, with input from SDP. DRM, DRS, MCM, JWR, and WB collected data. BJS performed analyses with input from DRM and TA. All authors contributed to the interpretation of the results. BJS wrote the first draft of the manuscript with DRM, TA, and SDP contributing to its revision prior to edits by all authors.

## Data accessibility statement

All data (DOI: 10.5281/zenodo.21402472) and code (DOI: 10.5281/zenodo.21403405) are archived through Zenodo. Code is also available on GitHub (https://github.com/bsmity13/elk_RT).

## Boxes

### Box 1: A graphical model of elk defensive traits by age and predator

Plastic defensive traits balance costs and benefits, and we hypothesize that the balance of that tradeoff should change with (1) prey senescence and (2) predator traits. We illustrate adaptive trait levels using a graphical framework based on ecological theory (Peacor et al. 2013). The framework makes our assumptions underlying key relationships and their effects explicit and allows us to deduce their logical consequences. We assume higher defensive trait expression incurs lower predation (benefit) and lower growth (cost). We conceptualize fitness as a net rate combining gains from growth and losses from predation. Here, “growth” describes all aspects of fitness other than mortality from the focal predator. The adaptive value of the defensive trait is the maximum of the fitness curve (yellow circles in Fig. 1); i.e., the point at which the marginal cost of increased defense exceeds its marginal benefit. The relationships between the defensive trait expression (x-axis), growth (blue lines), and predation (red lines) are based on the biology of our system: elk (*Cervus canadensis*) navigating predation risk from wolves (*Canis lupus*) and cougars (*Puma concolor*).

#### Defensive Trait

Elk movement and habitat selection behaviors are a defensive trait used to avoid risky places at risky times (see main text). Habitat selection reflects a tradeoff between food and safety: low values of the defensive trait emphasize food (risk-taking), whereas high values emphasize safety (risk avoidance).

#### Growth

We represent the growth rate vs. defensive trait relationship as a concave-down function, i.e., differences in risk avoidance have a larger effect at high risk-avoidance levels. Elk balance forage quantity with digestion (“handling”) time, consistent with a type-II functional response. As elk further express their defensive trait by reducing their foraging effort, their growth becomes less limited by handling and the curve becomes steeper.

##### Senescence effect

For prime-aged elk, moderate risk avoidance has a relatively small effect on growth (Fig. 1A, C) because individuals can satisfy their energetic needs with intermediate foraging effort. For senescent elk, greater foraging effort is required to meet their energetic requirements. Therefore, risk avoidance becomes more costly with age, causing the growth curve to decline more steeply (Fig. 1B, D).

#### Predation

We represent the relationship between predation rate and defensive trait expression as a concave-up function, such that increases in risk avoidance yield diminishing reductions in predation risk. This is consistent with proportional hazard reduction, where a unit increase in the defensive trait produces a percentage decrease in risk, as conventionally assumed in both survival (hazard rates) and habitat selection (relative selection strengths – see main text).

##### Senescence and predator hunting mode effects

Wolves (Fig. 1A-B) are coursing predators that target vulnerable individuals. For prime-aged elk, wolf predation risk is relatively low, represented by a curve with a small y-intercept (Fig. 1A). Senescent elk are much more vulnerable to wolves, resulting in a larger y-intercept (Fig. 1B). We assume similar decay rates across age classes, reflecting that risk avoidance reduces encounter and/or attack rates but cannot fully offset intrinsic vulnerability. Thus, even highly risk-avoidant senescent elk experience higher wolf predation than prime-aged elk.

Cougars are stalk-and-chase predators with specialized hunting adaptations that increase lethality upon encounter. Thus, predation risk is higher for cougars than for wolves across age classes, represented by higher intercepts (Fig. 1C-D). However, because cougars rely on specific stalking habitats, risk avoidance is more effective at reducing predation risk, represented by steeper declines in the predation curve (Fig. 1C-D). By contrast, wolves operate over broader habitat domains, making spatial risk avoidance less effective per unit trait change.

#### Predictions

The combined effects of age-dependent growth costs and predator-specific changes in vulnerability generate qualitative differences in optimal defensive trait expression with senescence (Fig. 1E). Senescence increases the cost of risk avoidance, such that equivalent levels of defense require larger benefits to remain adaptive. For wolves, risk avoidance has little benefit for prime-aged elk but becomes more beneficial as vulnerability increases with senescence.

Despite higher costs, the increased benefit can shift the fitness optimum toward greater risk avoidance. In contrast, for cougars, risk avoidance is already advantageous for prime-aged elk due to high lethality and the effectiveness of avoidance. With senescence, however, the increased cost of defense outweighs additional benefit, potentially leading to little change or even increased risk-taking (Fig. 1E).

These relationships generate two predictions: (1) senescence will tend to increase risk avoidance in response to coursing predators (e.g., wolves) but not necessarily stalk-and-chase predators (e.g., cougars), and (2) predator-specific differences in how behavior reduces risk will determine the direction and magnitude of age-related changes in defensive traits.

**Figure 1.**
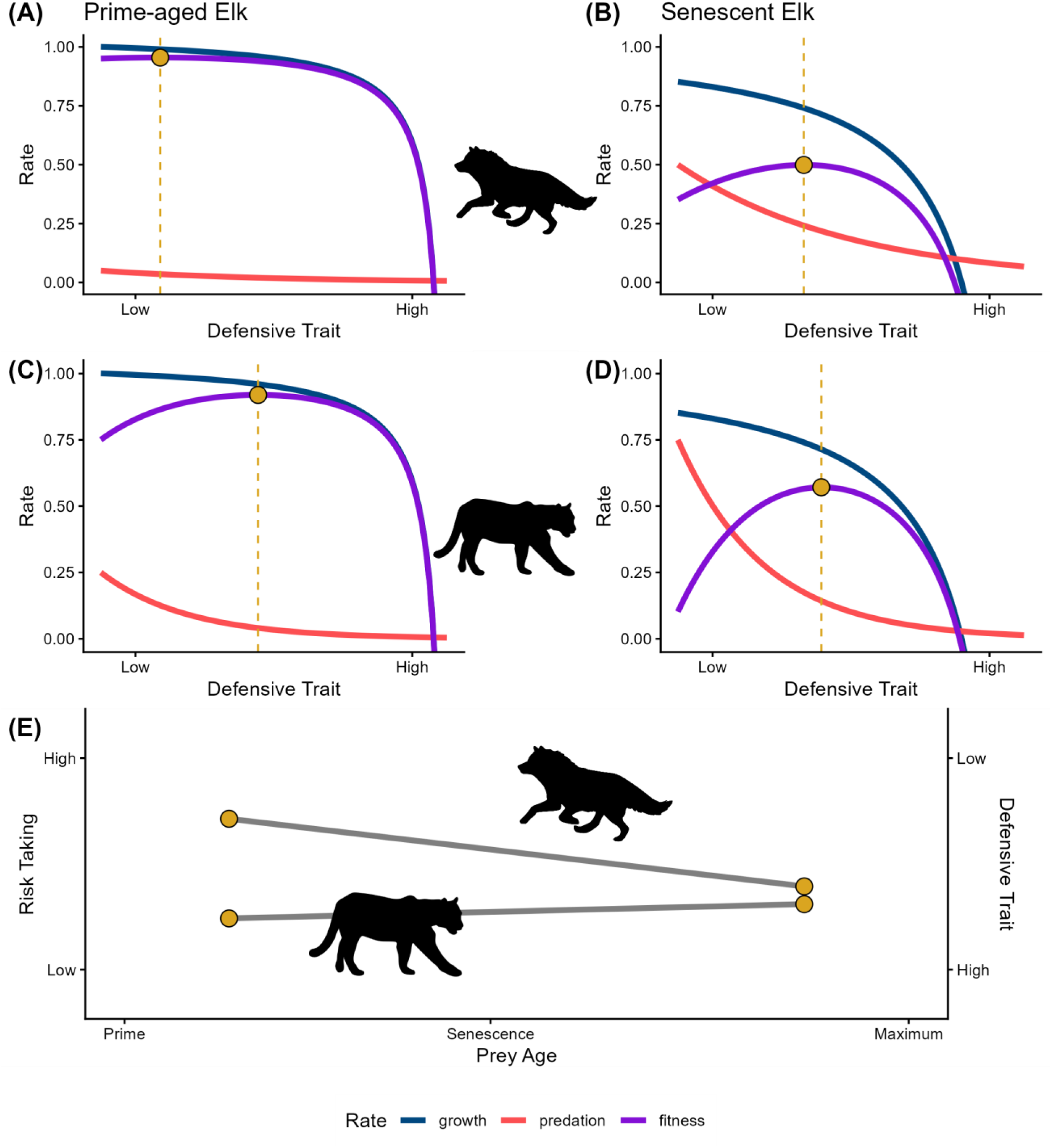
A graphical model of how predator hunting mode affects defensive investment of senescing prey, based on Peacor et al. (2013). Fitness (purple) is conceptualized as growth rate (blue) minus predation rate (red), where “growth” refers to all components of fitness not related to predation by the focal predator. Yellow circles show the value of the defensive trait that maximizes fitness. Cougars are more lethal than wolves [greater Pr(Kill|Attack)], and both are expected to be more lethal when attacking old vs prime-aged elk, but this age effect is expected to be significantly larger for wolves (who chase their prey) than for cougars (who stalk their prey). (A) Prime-aged elk and wolves. (B) Senescent elk and wolves. (C) Prime-aged elk and cougars. (D) Senescent elk and cougars. (E) Predicted change in the defensive trait with age with respect to the two predators. We refer to high values of the defensive trait as “risk avoidance” and low values as “risk-taking.”

## Appendix S1: Estimating predation risk in space

### Summary

We modeled spatial variation in predation risk (separately for wolves and cougars) from a dataset of elk kills (cows and calves only) attributed to VHF- or GPS-collared predators. We used negative binomial generalized additive models (GAMs) to estimate the relationship between landscape characteristics (canopy openness, terrain roughness, snow-water equivalent) and expected kills, while controlling for (1) elk density, (2) collared predator space use [the observation process], and (3) random effects in time and space.

We used the fitted wolf/cougar models to predict the distribution of wolf-/cougar-killed elk, conditional that (1) elk were distributed uniformly and (2) observed predators were distributed uniformly; the purpose of this prediction was to estimate spatial risk independent of elk distribution – which we assume includes some signal of risk avoidance (see Smith et al. 2023) – and the observation process, i.e., the location of collared predators that generated the dataset. We used this prediction as our estimate of predation risk.

We made separate predictions for early winter (November 15 – December 15) and late winter (March) seasons, which we used in the main text to estimate the effect of risk on hourly elk behavioral responses.

### Background

Our primary goal in this analysis was to estimate spatial variation in predation risk. We defined predation risk as the probability of an elk kill, given the location; this is the probability of being killed in a place and *not* necessarily what the elk perceive as risky. A secondary goal of this analysis was to interpret the ecological meaning of the relationships between the spatial predictors of risk and the expected number of kills.

### Methods

#### Data Collection

Wolves and cougars were monitored during two intensive periods each winter: Early Winter (November 15 – December 15) and Late Winter (March 1 – March 31). During each of these periods, telemetered predators were monitored for evidence of a potential kill, and all potential kill sites were visited to determine the attributes of the prey species. We restricted the data used here to only cow or calf elk to reflect risk to a female elk or her offspring. The winter distribution of cow/calf kills differs from the winter distribution of bull elk kills (Kohl *et al*. 2018), which likely reflects spatial segregation of cows/calves from bulls during winter (Houston 1982). Note that all GPS-collared elk considered here were adult females. We included data from wolves monitored from 2001 – 2020 and cougars monitored from 2016 – 2020. We refer to the years by the “winter” in which they occurred, defined by the calendar year of that January. For example, winter 2020 equaled “Early Winter 2020” (November 15, 2019 – December 15, 2019) and “Late Winter 2020” (March 1, 2020 – March 31, 2020). We included data from 40 winter-seasons for wolves and 9 winter-seasons for cougars.

We estimated predation risk using this large dataset of kills and generated predicted risk surfaces for 2016 – 2020 for use in this analysis. These data emerge via two processes: (1) a state process whereby predators are killing elk on the landscape, and (2) an observation process by which only the monitored predator’s kills are observed. We sought to control for the observation process while estimating the underlying state process.

#### Data Preparation

We estimated predation risk in discrete space by rasterizing the northern range study area with 250 m x 250 m pixels. We used this same raster to represent all data, including the kills and all potential predictors of interest. For each winter-season and for each predator species, we compiled the following datasets: (1) predator kills, (2) elk density, (3) predator space-use intensity, (4) number of overlapping wolf packs, (5) snow-water equivalent (SWE), and (6) canopy openness and terrain roughness.

We represented predator kills as the total number of elk kills (cows and calves only) within that pixel in a given winter-season. We did not consider kills of alternative prey items (including bull elk).

We estimated elk density using predictions from the model of Smith et al. (2023). That elk density model predicts expected elk density on a raster of 1-km pixels, while considering density-dependent changes in habitat selection by elk. This predictor is thus model-based, but recall that virtually all remotely sensed predictors available at a landscape scale are also necessarily estimates from a mechanistic or phenomenological model. We estimated elk density for each winter-season by passing the density model all necessary covariates, including the corresponding SWE for that winter-season. That is, SWE is the only covariate in the elk density model that differed between early and late winter within a single year. We resampled the estimated density to our 250-m pixels using a nearest neighbor algorithm, and then divided the total by 16 (because there are 16 250-m pixels in a 1-km pixel) to retain the correct total population size for that winter.

We estimated predator space-use intensity by estimating a utilization distribution (UD). Two challenges arise in estimating this distribution: (1) predators were tracked with varying temporal resolution in different years, frequently enough in recent years that non-independence of locations becomes a concern; and (2) in many instances, multiple wolves in the same pack were tracked, another potential source of non-independence. While methods exist that can estimate a UD while accounting for temporal autocorrelation in location data (e.g., AKDE; Fleming et al. 2015), those methods are individual-based and do not estimate social interactions of pack members. Instead, we chose to randomly sample one location/pack/day and treat the sampled locations as independent. We then used an ordinary kernel density estimator (KDE) to estimate the UD. For consistency, we followed the same procedure for cougars, randomly sampling one location/cougar/day. We estimated the UD, specific to each predator, on our 250-m raster. We truncated each KDE at the 99^th^ isopleth (the smallest polygon containing a cumulative 99% probability of use), renormalized so that the values summed to 1, and multiplied by the group size (the pack size for wolves, or 1 for cougars [including only adults that hunt prey]).

Thus, the resulting raster was interpreted as the expected number of predators (either wolves or cougars) in that pixel at any given point in time, and we refer to this term as “predator density” (specifically, “wolf density” or “cougar density”). For fitting each predator model, we retained only those pixels where predator density was > 0, to avoid including pixels where no predator was being monitored in that winter-season.

In addition to predator intensity of use, we also estimated the number of overlapping wolf packs in a given pixel. Previous work suggests that territoriality in wolves lowers predation risk where territories overlap (Kauffman *et al*. 2007), so we sought to include number of overlapping packs to account for this. While monitoring of wolf packs within Yellowstone National Park is nearly exhaustive, with telemetry collars in most packs during most years, cougar monitoring is not designed to be as exhaustive. Therefore, we did not include a similar estimate of overlap for cougars, because we did not assume we knew the location of most cougars in the system with confidence. Note also that female cougars often have overlapping home ranges, whereas males are much more territorial.

We estimated snow-water equivalent in each pixel with the SWE product from Daymet 4.0 (Thornton et al. 2022). SWE has been shown to be an important predictor of elk use (Mao et al. 2005; Smith et al. 2023) and predation risk (Kauffman et al. 2007) in this system. Daymet provides daily estimates of SWE on a 1-km raster. To estimate snow for a given winter-season, we downloaded all days of SWE data from Daymet, took the mean in each pixel, and then resampled to our 250-m raster using a nearest neighbor algorithm.

Canopy openness (hereafter, “openness”) and terrain roughness (hereafter, “roughness”) together have been shown to be important predictors of elk predation risk in this system (Kohl et al. 2018, 2019; Smith et al. 2023). We estimated openness using the cover product from the Rangeland Analysis Platform (RAP; Jones et al. 2018), which has a native resolution of 30-m.

RAP estimates the percent cover of trees, and we calculated percent openness as 100 – tree cover, then we resampled to our 250-m raster using bilinear interpolation. RAP cover data were estimated annually, and we used the data from the year prior to each winter of interest (e.g., for winter 2020, we used RAP’s tree cover estimate for 2019). To estimate roughness, we obtained a digital elevation model (DEM) from the National Map’s 3DEP program (U.S. Geological Survey 2020), projected to a 30-m grid. We calculated roughness in R using the `terrain()` function from the `raster` package with `opt = "roughness"` (Hijmans 2022). We then resampled the roughness raster to our 250-m raster using bilinear interpolation.

#### Modeling

We fit a Generalized Additive Model (GAM) for each predator species to estimate the expected number of kills in each pixel. GAMs use penalized splines to represent smooth predictors, thus the relationship between a predictor and the response variable can take a non-linear form that is not pre-specified by the user, and cubic regression splines (and their generalizations) used in GAMs can theoretically represent any smooth function (Wood 2017). This is useful in this context because, for example, the expected relationship between prey density and kill rate (the functional response), is typically expected to be non-linear in most systems; an arbitrary smooth function can capture this without specifying the functional form of that relationship parametrically. The penalty (as in “penalized splines”) reduces the total degrees of freedom used by each smooth term; i.e., the data inform how much complexity a spline should have, and as long as the user provides enough basis functions a priori, the model will select an appropriate number of basis functions to parsimoniously represent the relationship. The penalty induces shrinkage exactly analogous to the shrinkage of random effects in a generalized linear mixed model (Wood 2017). For that reason, GAMs can also be used to flexibly fit random effects and even spatial random effects (i.e., a Gaussian process term, a.k.a., kriging).

The two predator models had nearly the exact same predictors (Table S1.1). Both models included these four smooth (penalized spline) terms: (1) the natural logarithm of elk density, (2) the natural logarithm of predator density, (3) SWE, and (4) a tensor smooth of openness and roughness (analogous to an interaction between the two variables). Rationale for the log-transformations of elk density and predator density is provided below. Both models included two random effects: (1) a random intercept for winter and (2) a Gaussian process to account for unmodeled spatial autocorrelation (a spatial random intercept). We used a Matern covariance function for the Gaussian process, as suggested by Kammann and Wand (2003) and recommended by Wood (2017). Both models also included a fixed effect of season, allowing the average number of kills to differ between early and late winter. The wolf model additionally included a smooth term for the number of wolf packs (described above).

**Table S1.1.** Predictors in predation risk models. We modeled the risk to elk from wolves and cougars using two separate models. The models were nearly identical, except that the wolf model accounted for the number of overlapping territories, whereas the cougar model did not. All “smooth” terms, including the tensor smooth, used cubic regression spline basis functions.

| **Variable** | **Form** | **Wolf**<br>**Model** | **Cougar**<br>**Model** | **Predicted Effect** |
| --- | --- | --- | --- | --- |
| log Elk Density | Smooth | X | X | Increasing, saturating |
| log Predator Density | Smooth | X | X | Increasing, saturating |
| Wolf Packs | Smooth | X |  | Monotonically decreasing |
| SWE | Smooth | X | X | Monotonically increasing |
| Openness, Roughness | Tensor smooth | X | X | High at high openness and low roughness for wolves; high at intermediate openness and high roughness for cougars |
| Spatial RE | Gaussian<br>process | X | X | No prediction |
| Winter RE | Random<br>intercept | X | X | No expectation |
| Season | Parametric<br>fixed effect | X | X | Increased risk in late winter |

The response variable for both models was the count of kills in each pixel, and we could model the counts as either a Poisson or a negative binomial random variable. The canonical link function for either would be the natural logarithm. We fit the full model for each predator, assuming a Poisson random variable. We used the R package DHARMa to evaluate the model based on simulated residuals, checking for zero inflation, overdispersion, uniformity of residuals, outliers, and heteroskedasticity. The wolf model passed all of our diagnostic checks, whereas the cougar model showed evidence of overdispersion. We refit the cougar model, treating the count of kills as a negative binomial random variable, and the resulting model passed all diagnostic checks. Thus, our final models were a Poisson GAM for wolf risk and a negative binomial GAM for cougar risk.

#### Log-transformation of densities

The Poisson/negative binomial GAMs used the natural logarithm (hereafter, “log”) as the link function. Thus, the linear predictor represented the mean of the Poisson/negative binomial distribution on the log scale. Therefore, we log-transformed the elk and predator densities to match the scale of the response variable; in other words, we log-transformed the elk and predator densities to match the log-transformed kills. To illustrate why, we will use the analogy of a generalized linear model (GLM) by omitting any random effects and reducing the smooth functions of the GAM to simple linear terms. Briefly, the linear predictor for expected elk kills in spatial cell *i*, would be:

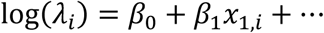

for each of the predictors (*x*) in the model. The units of *λ_i_* are expected number of kills. If we hypothesized that the expected number of kills scaled linearly with elk abundance and predator abundance, we could include the log of these terms as an offset, a predictor whose coefficient is assumed to be 1. In that case:

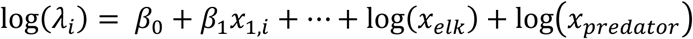

By exponentiating both sides (“back-transforming” the prediction), we would get:

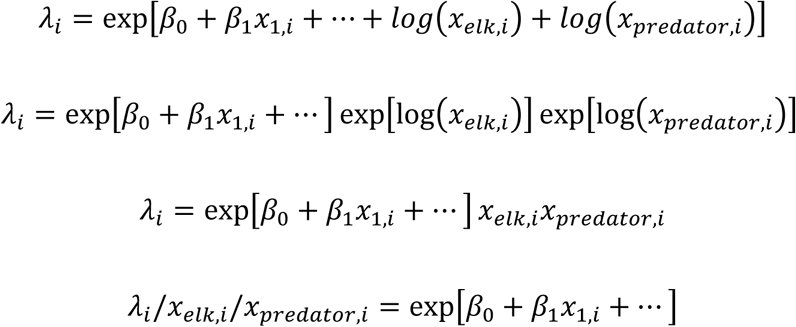

Thus, the predictors remaining on the right-hand side of the equation give the expected kills/elk/predator, since the exponentiation of the right-hand side canceled the log transformation of elk and predator densities. In the case of the GAM, we are implying that we do not assume the relationship between kills and elk/predator abundance is linear (we will estimate the functional form), but we use the log-transformation so that the predictor is linear when back-transformed. If we predict expected kills (*λ_i_*) from the fitted GAM with (log) elk density and (log) predator density held constant (see below), the expected number of kills/elk/predator in any pixel is linearly proportional to *λ_i_* by a constant given by evaluating the estimated smooth functions at the constant value of elk and predator densities.

#### Predicting Predation Risk

The primary goal for our fitted models was to predict spatial variation in predation risk to elk. We limited the spatial domain of our prediction to the portion of the northern range inside of Yellowstone National Park. We generated a prediction for each predator for each winter-season. We used winter-season-specific risk layers to estimate how elk respond to predation risk at a fine temporal scale (1 hour).

We conceptualized risky places as those places where an elk would be most likely to be killed, conditioned on: (1) a constant number of elk present and (2) a constant number of predators present. The rationale for (1) is that if elk tend to avoid risky places, there will be very few elk in the riskiest places and thus fewer kills than expected based on risk alone; we want our prediction to remove the signal of variable elk density in the kills. The rationale for (2) is twofold: (a) more predators will tend to make more kills, and (b) monitoring of telemetered predators is the observation process that underlies the dataset. Thus, we created predictions where we held both elk density and predator density constant at the mean observed value for each winter-season. For the wolf model, we also held the number of wolf packs constant at 1.

#### Interpreting Relationships

The secondary goal for our fitted models was to interpret the relationship between the smoothed covariates and the kills. We developed a priori qualitative expectations for each of the five primary smooth terms: (1) elk density, (2) predator density, (3) wolf packs, (4) SWE, and (5) openness x roughness.

We expected that the smooth term for elk density was proportional to the predator’s functional response. The functional response describes how kills/predator/time changes with prey density. In our model, predicted kills (per season, a unit of time) are proportional to kills/elk/predator (as explained in *Log-transformation of densities*); thus, we interpreted the fitted smooth as proportional to the functional response. We expected a type II functional response for both predators, i.e., monotonically increasing, but saturating. It was possible that a Type III functional response operated in this system, but detecting the difference between Type II and Type III from noisy observational data is difficult (Metz *et al*. 2020).

We expected the smooth term for predator density would also be monotonically increasing and saturating. We interpreted this effect as a predator-dependent functional response. The predator-dependent functional response describes how kills/predator/time change with predator density, but in many predator-prey models, it is expected to be a linear function.

We expected the smooth term for wolf packs would be monotonically decreasing due to territoriality.

We expected the smooth term for SWE would be monotonically increasing for wolves and cougars, because elk are more vulnerable to predation in deep snow.

Prior research on the effect of openness and roughness guided our expectations for this term (Kohl et al. 2019). We expected wolf kills to be greatest in pixels with high openness and low roughness, whereas we expected cougar kills to be greatest in pixels with intermediate openness and high roughness.

#### Model Validation

We randomly selected some winter-seasons of predator data to withhold for validation. For wolves, we had 20 early winters and 20 late winters, and we randomly selected 3 of each to withhold for testing. For cougars, we only had 4 early winters and 4 late winters, so we randomly selected 1 of each to withhold for testing.

We performed two different validation procedures on each model to test different aspects of their predictive ability, and both were based on simulation of new datasets from the fitted model. First, we calculated the expected number of total kills in each of the validation winter-seasons. We compared that with the observed number of kills in the testing data by looking for overlap with the predicted confidence intervals (considering parametric uncertainty) and prediction intervals (considering parametric uncertainty and Poisson/negative binomial variance). Second, we used the Boyce index popular for validating species distribution/habitat selection models to evaluate how well the model predicted the location of kills in geographic space (Boyce *et al*. 2002; Wiens *et al*. 2008). The Boyce index uses Spearman’s correlation coefficient to quantify the agreement between model predictions and observed counts, so ranges between -1 and 1.

### Results

#### Model Evaluation and Validation

Both the wolf and cougar models passed all diagnostics and showed no signs of zero-inflation, overdispersion, or other issues in the residuals. All smooth terms had effective degrees of freedom (EDF) much less than the number of knots, indicating a sufficient number of knots. The fitted wolf model explained 19.1% of the deviance in the wolf kill data, and the fitted cougar model explained 41.9% of the deviance in the cougar kill data. While the covariates did not explain the majority of the deviance in the models, they still performed well under cross-validation (see below), indicating they were reliable for inference.

The models predicted the observed number of total kills in 5 out of 6 validation winter-seasons for wolves (Fig. S1.1A) and 2 out of 2 validation winter-seasons for cougars (Fig. S1.1B). The Boyce index averaged 0.80 (range = 0.58 – 0.89) across all validation winter-seasons for wolves and averaged 0.73 (range = 0.62 – 0.84) across all validation winter-seasons for cougars. We concluded that the predation risk models had good predictive ability and proceeded to use them to predict predation risk across space.

**Figure S1.1.**
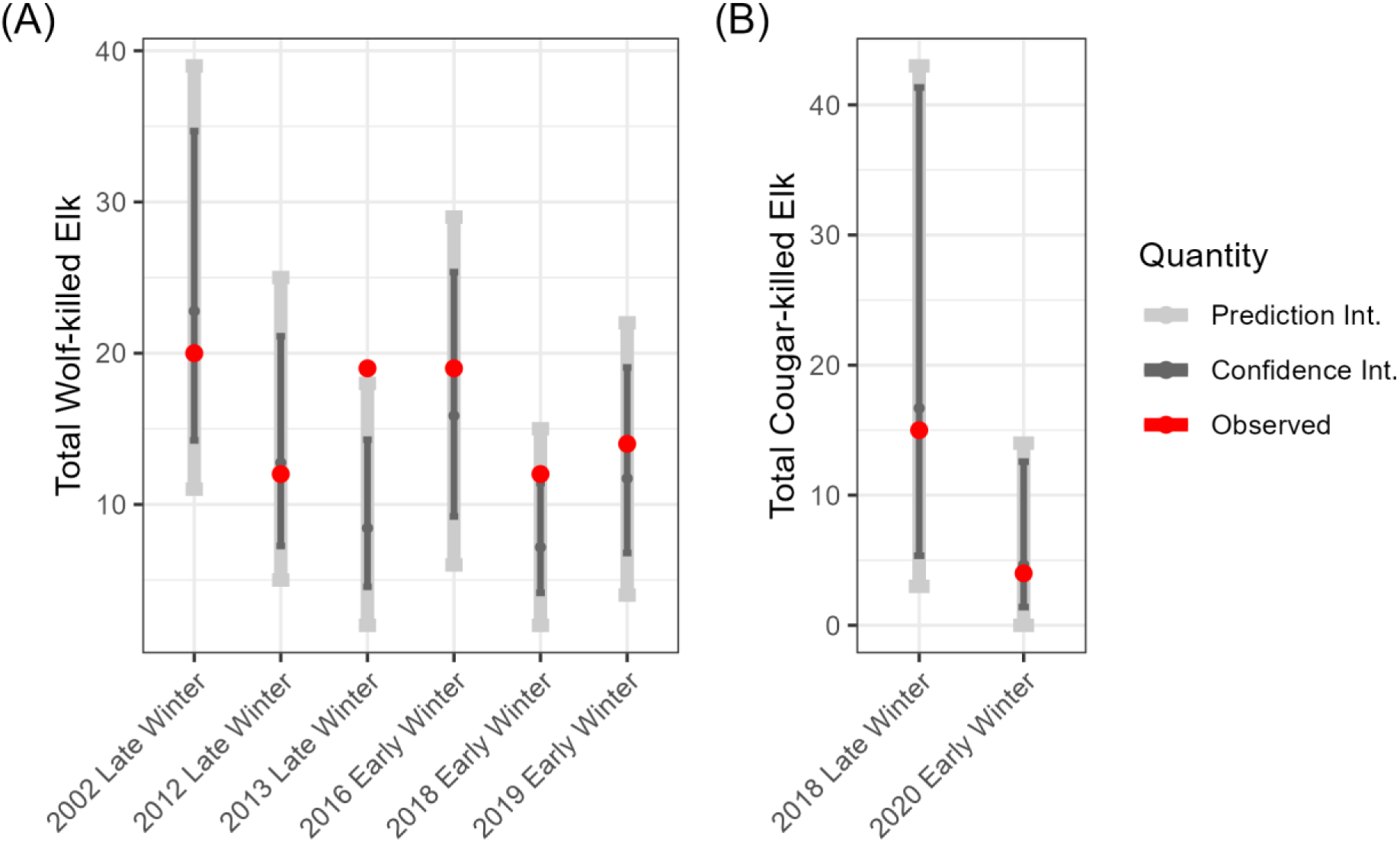
Model validation. We predicted total elk kills for each out-of-sample winter-season of data for both predator models. We considered the predictions to be correct if the observed value fell within the prediction interval. The wolf model predicted the observed number of kills in 5 out of 6 years (A), and the cougar model predicted the number of kills in 2 out of 2 years (B).

#### Interpreting Relationships

#### Risk from Wolves

The elk density curve was monotonically increasing and saturating, as we predicted. The resulting shape matches a type II functional response (Fig. S1.2A). This is partially due to the log-transformation, since in the absence of strong pattern, penalized splines approximate a straight line, which in this case is monotonically increasing due to the log-transform. This spline could have taken on another shape (e.g., Type I or Type III functional response) had the data included a strong signal of this. The wolf density curve was also monotonically increasing and saturating, matching our predictions, but the magnitude of the effect was much larger and with less uncertainty (Fig. S1.2B). The wolf pack curve was generally monotonically decreasing, except for a slight bump at 3 packs, as we predicted (Fig. S1.2C). This pattern agrees with the findings of Kauffman et al. (2007). The SWE curve was generally increasing, but with large confidence bounds (Fig. S1.2D). The magnitude of the predicted effect is large at high values of SWE, but the uncertainty remains large at these values, as well. Perhaps this was due to the low resolution of the SWE data (1-km pixels) obscuring details on the landscape, or perhaps the effect was weaker due to averaging the daily SWEs across each month-long season. The tensor for openness and roughness also matched our predictions, with the greatest contribution to kills coming in places that have high openness (positive effect > ∼80%) and low roughness (positive effect < ∼ 30 m; Fig. S1.2E).

**Figure S1.2.**
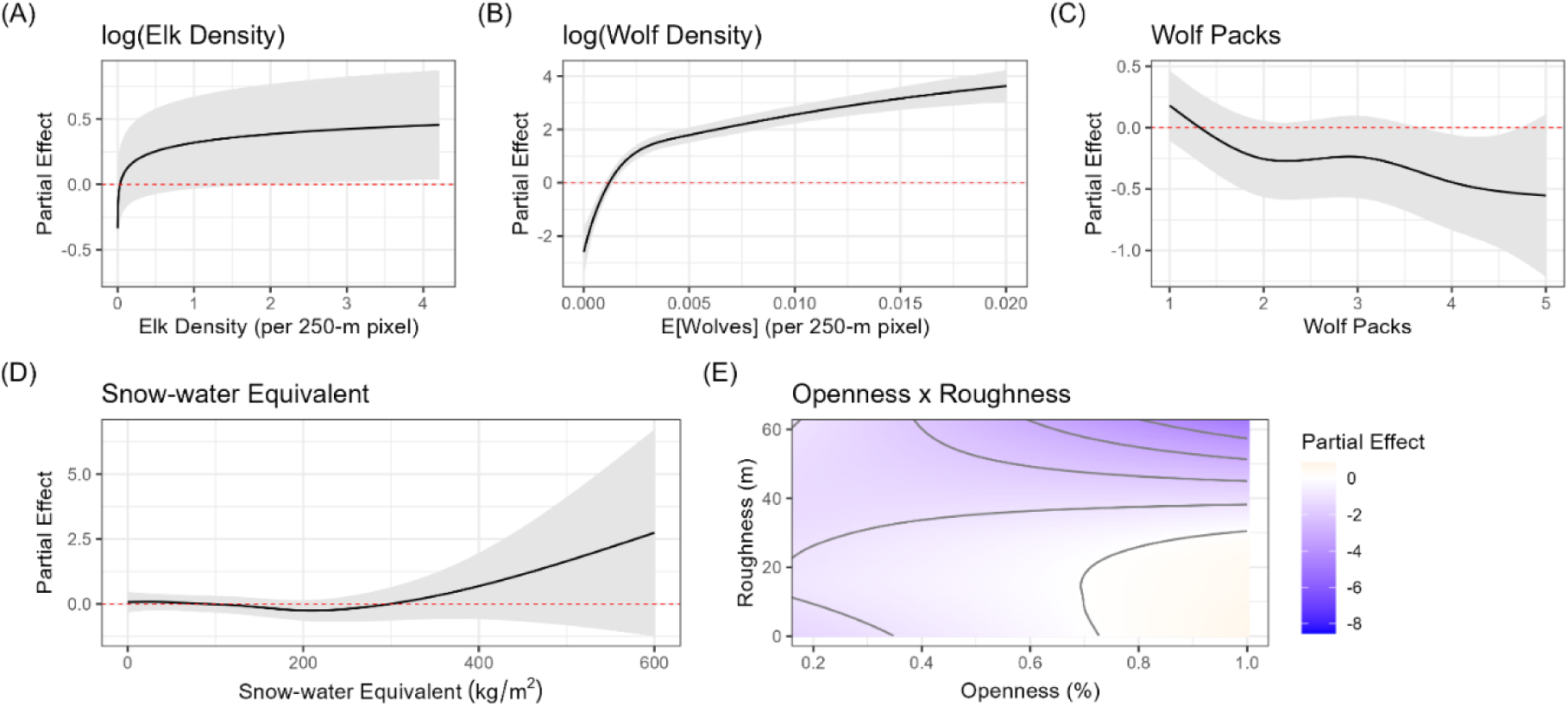
Wolf model smooth terms. The partial effects (y-axis) describe the contribution of each term to the linear predictor of the model. They can be used to compare relative effect sizes and note that the scale of the y-axes differs across panels. (A) The effect of elk density. (B) The effect of wolf density. (C) The effect of overlapping wolf packs. (D) The effect of snow-water equivalent. (E) The effect of the tensor for openness and roughness.

The random intercept for winter had a generally small effect (Fig. S1.3A). Most winters were not significantly different from 0 (i.e., the baseline number of kills equaled the long-term average), but there were a few exceptions. The winters of 2005 and 2020 had fewer wolf kills than average, while the winter of 2011 had more wolf kills than average. The fixed effect of season was significant in the wolf model (p = 0.001). The estimated coefficient was 0.48, meaning that there were on average exp(0.48) = 1.6 times more kills in Late Winter than in Early Winter. The Gaussian process term (the residual spatial autocorrelation) tended to increase the number of kills along the middle of the study area, following the Yellowstone River (Fig. S1.3B). That is, the model predicted kills in this area (orange pixels in Fig. S1.3B) were higher than expected based on the other covariates alone.

**Figure S1.3.**
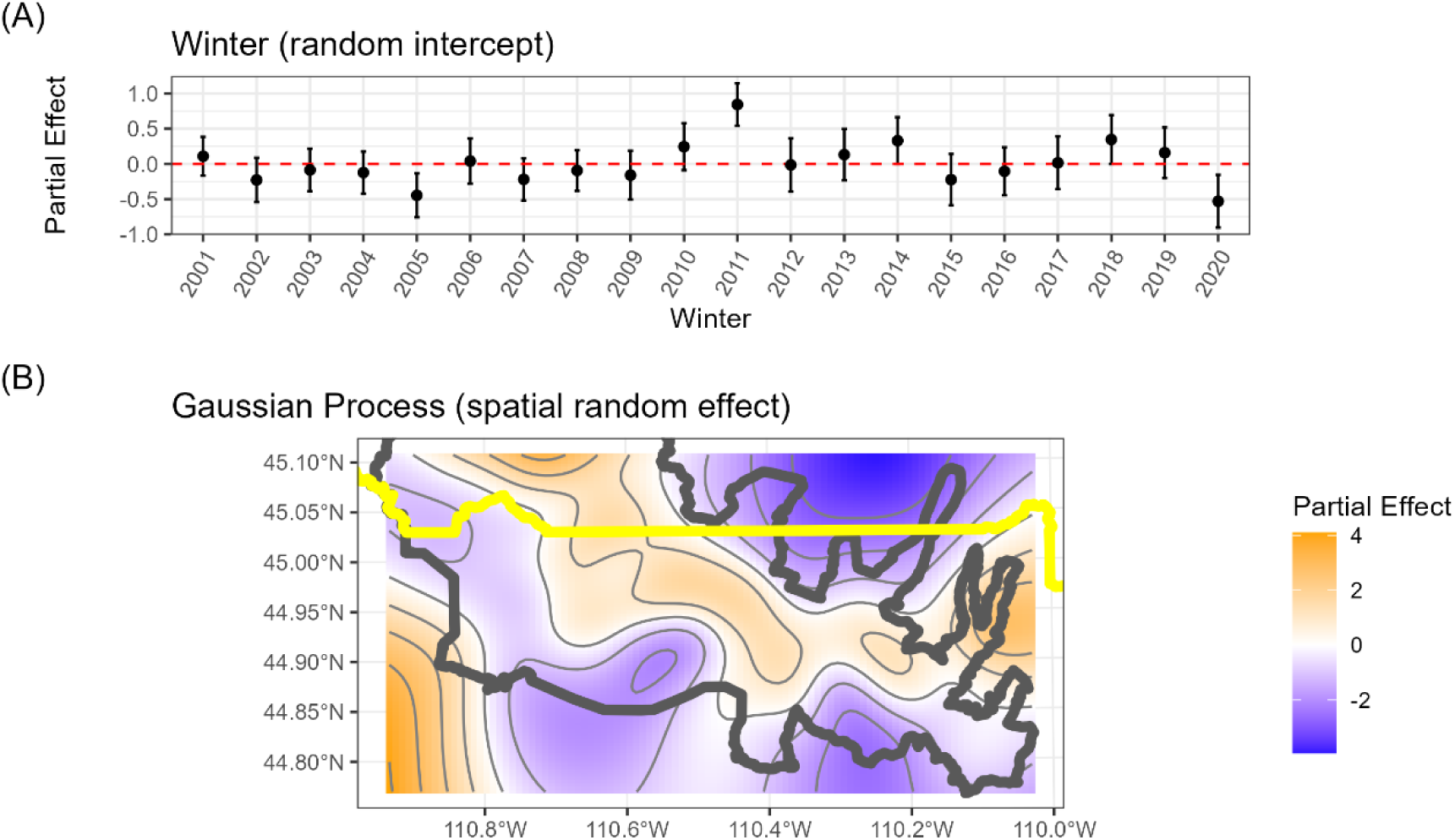
Wolf model random effects. The partial effects (y-axis) describe the contribution of each term to the linear predictor of the model. They can be used to compare relative effect sizes, and note that the scale of the y-axes differ across panels. (A) The effect of winter, included as a random intercept. (B) The spatial random effect, modeled as a Gaussian process (a.k.a., krigging).

#### Risk from Cougars

The elk density curve was essentially flat when considering the large confidence intervals, contrary to our predictions (Fig. S1.4A). The cougar density curve was monotonically increasing and saturating, matching our predictions, and the effect was the largest of any term (Fig. S1.4B). The SWE curve was linearly decreasing, but with large confidence bounds, essentially indicating no effect (Fig. S1.4C). Perhaps this is due to the low spatial or temporal resolution of the SWE data, as we mentioned for the wolf SWE effect. The tensor for openness and roughness also matched our predictions, with the greatest contribution to kills coming in places that have intermediate to high openness (positive effect from ∼ 45 – 100%) and high roughness (positive effect > ∼ 20 m; Fig. S1.4D). Compared with wolves, the importance of roughness is greater for cougars. Compared with previous research in this system, it also suggests more cougar kills in the open than previous research has indicated (Kohl *et al*. 2019), mainly in rough terrain.

**Figure S1.4.**
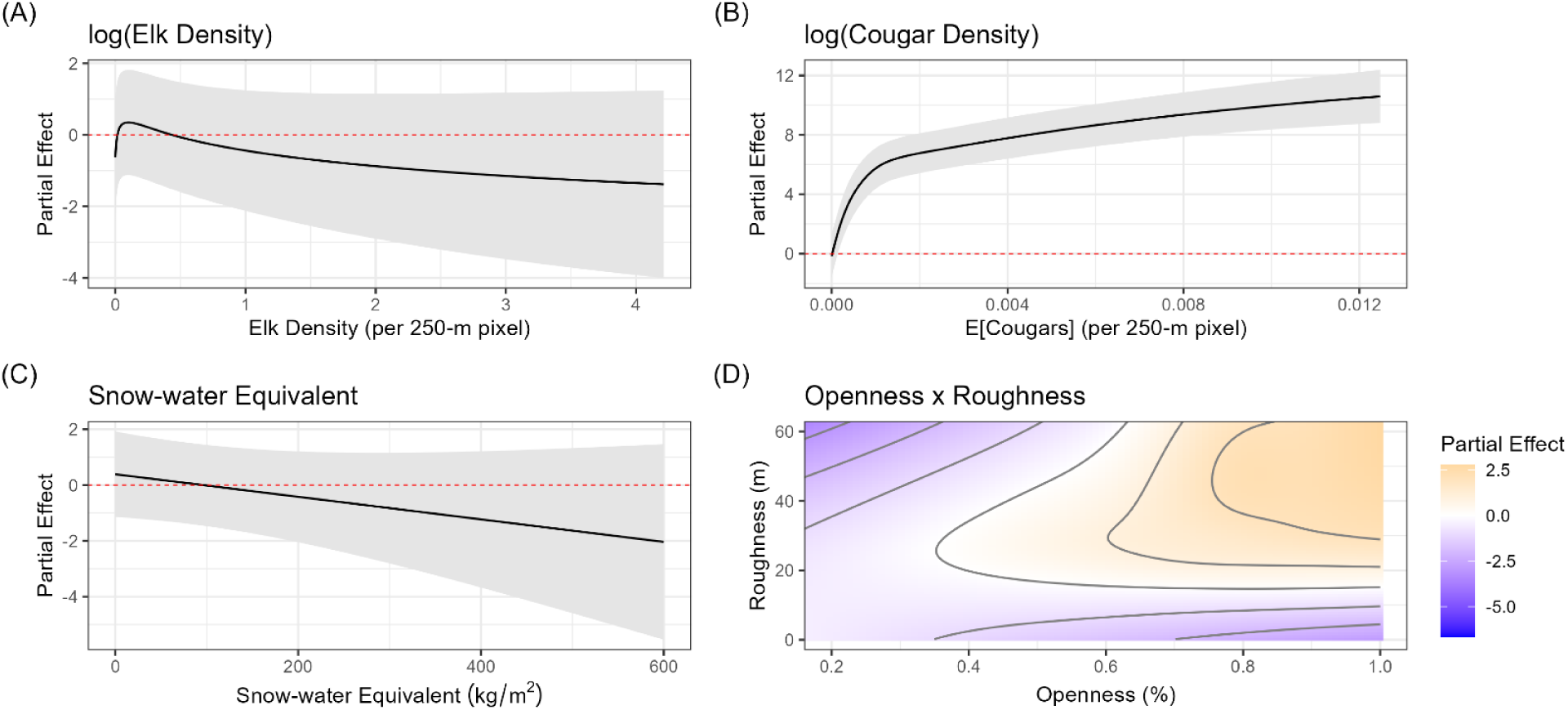
Cougar model smooth terms. The partial effects (y-axis) describe the contribution of each term to the linear predictor of the model. They can be used to compare relative effect sizes, and note that the scale of the y-axes differs across panels. (A) The effect of elk density. (B) The effect of cougar density. (C) The effect of snow-water equivalent. (D) The effect of the tensor for openness and roughness.

The random intercept for winter had virtually no effect (Fig. S1.5A), with no winters significantly different from 0 (i.e., the baseline number of kills was equal to the long-term average). The fixed effect for season was also not significantly different from 0 (p = 0.297). The Gaussian process term (the spatial random effect) was qualitatively different from that of the wolves, tending to increase kills in the rough terrain coming down from the interior of Yellowstone National Park (Fig. S1.5B).

**Figure S1.5.**
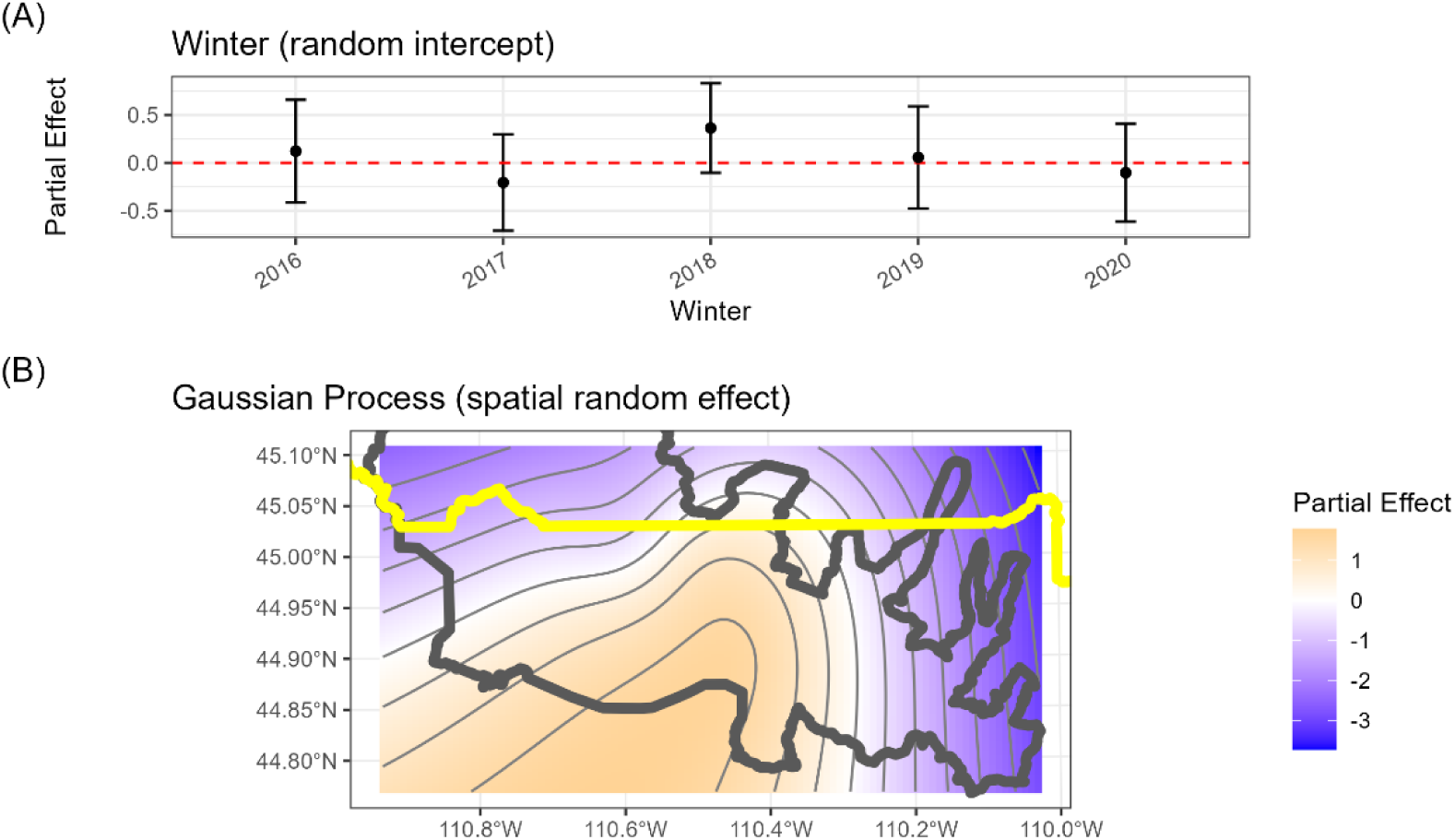
Cougar model random effects. The partial effects (y-axis) describe the contribution of each term to the linear predictor of the model. They can be used to compare relative effect sizes, and note that the scale of the y-axes differ across panels. (A) The effect of winter, included as a random intercept. (B) The spatial random effect, modeled as a Gaussian process (a.k.a., krigging).

#### Predicting Predation Risk

We will illustrate predicted predation risk with an example winter-season, “2018 Late Winter,” arbitrarily chosen, but recall that we produced specific predictions for each winter-season. For each predator, we presented four different panels: (A) expected kills under the full dataset, i.e., when all pixels of the raster have their observed value (from “2018 Late Winter”); (B) expected kills when elk are distributed evenly, i.e., when all pixels of the raster have their observed value, except that elk density is held uniform across the raster (at 1/*N_i_*, where *N_i_* is the total elk observed in that winter-season); (C) expected kills when the predator is distributed evenly, i.e., when all pixels of the raster have their observed value, except that predator density is held uniform across the raster (at 1/*P_i_*, where *P_i_* is the total predators observed in that winter-season) – for wolves, wolf packs were also held constant at 1; and (D) expected kills when both the elk and the predator are distributed evenly (a combination of B & C). We considered panel A to be the best prediction of where we expected to find kills, given the observation process (the red stars in panel A), while we considered panel D to be the best prediction of the places that are risky (i.e., the state process).

#### Risk from Wolves

The wolf model predicted the most expected kills to be concentrated near the center of our study area (Fig. S1.6A). Evenly distributing elk made little qualitative difference in the predictions (Fig. S1.6B), but evenly distributing wolves made a much larger difference in the distribution of expected kills (Fig. S1.6C). Thus, we observed a strong signal of the observation process. Accounting for both elk density and wolf density, our final prediction of predation risk (Fig. S1.6D) shows risk is distributed much more widely across the landscape that would be predicted based on the locations of the kills alone (red stars in Fig. S1.6A). We used Panel D as our measure of wolf risk.

**Figure S1.6.**
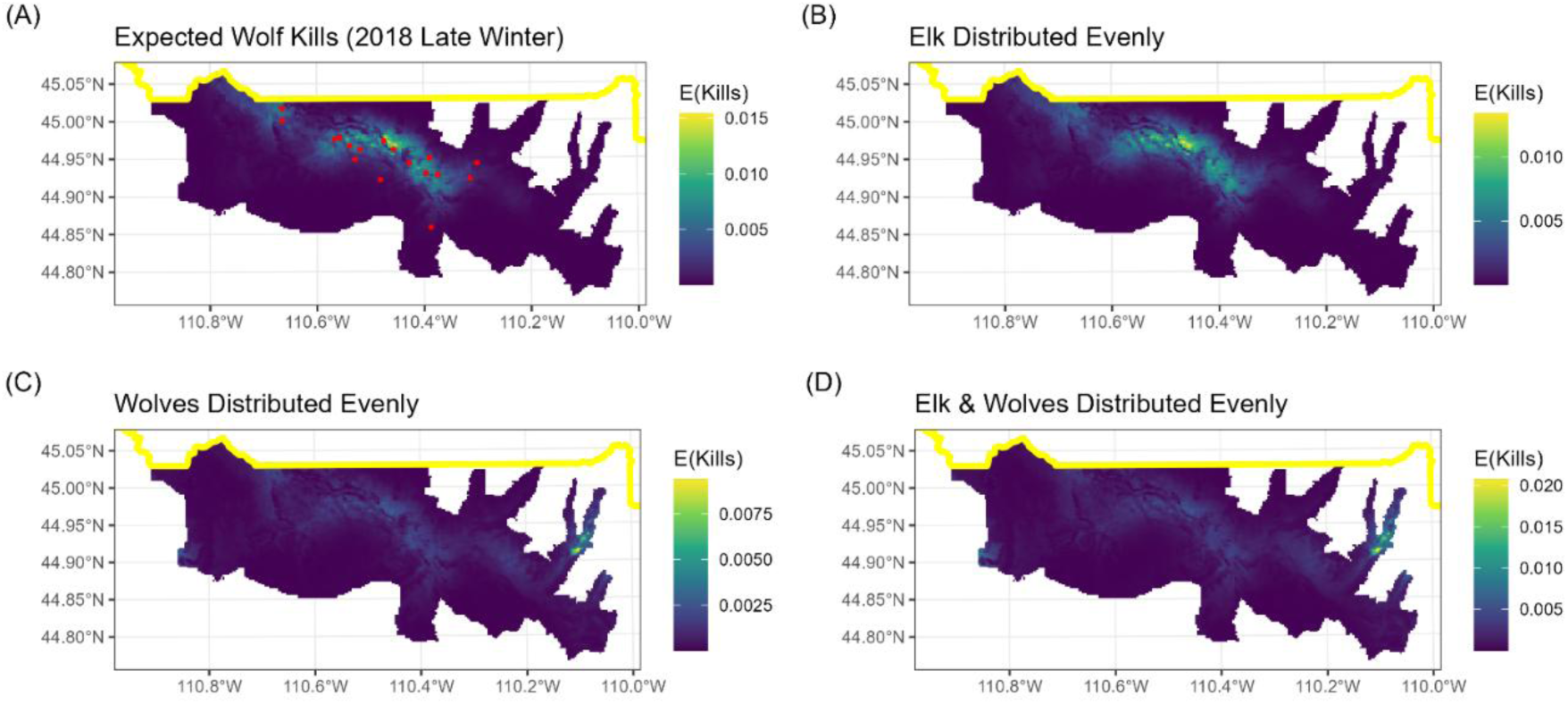
Wolf predation risk. We used our fitted wolf model to make four predictions (each panel) for each winter-season. Here, “2018 Late Winter” is used as an example. (A) Expected wolf kills given observed conditions. (B) Expected wolf kills given elk density held constant. (C) Expected wolf kills given wolf density (and number of packs) held constant. (D) Expected wolf kills given elk density and wolf density/packs held constant. The red stars in panel A are the observed location of wolf-killed cow or calf elk in late winter 2018 (N = 18 kills).

#### Risk from Cougars

Similar to the wolf model, the cougar model predicted the most expected kills to be concentrated near the center of our study area (Fig. S1.7A); however, expected kills were concentrated even more densely due to the smaller proportion of the cougar population monitored relative to the proportion of the wolf population monitored. Evenly distributing elk made little qualitative difference in the predictions (Fig. S1.7B), but evenly distributing cougars made a much larger difference in the distribution of expected kills (Fig. S1.7C). Thus, we observed a strong signal of the observation process. Accounting for both elk density and cougar density, our final prediction of predation risk (Fig. S1.7D) shows risk is distributed much more widely across the landscape than would be predicted based on the locations of the kills alone (red stars in Fig. S1.7A). We used Panel D as our measure of cougar risk.

**Figure S1.7.**
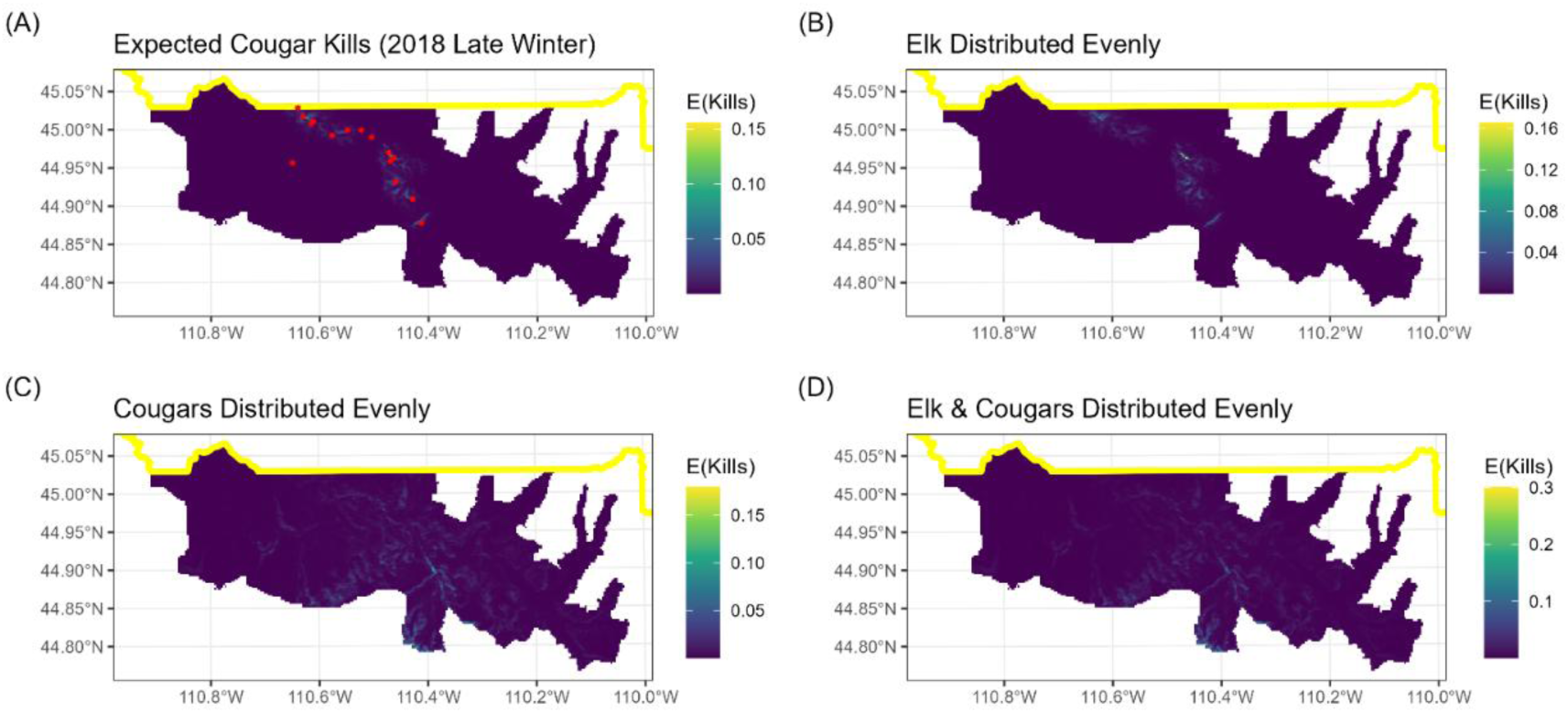
Cougar predation risk. We used our fitted cougar model to make four predictions (each panel) for each winter-season. Here, “2018 Late Winter” is used as an example. (A) Expected cougar kills given observed conditions. (B) Expected cougar kills given elk density held constant. (C) Expected cougar kills given cougar density held constant. (D) Expected cougar kills given elk density and cougar density held constant. The red stars in panel A are the observed location of cougar-killed cow or calf elk in late winter 2018 (N = 19 kills).

## Appendix S2: Estimating predation risk in time

### Summary

We modeled temporal variation in predation risk by modeling predator activity patterns (wolves and cougars separately) as a function of time of day during early winter (November 15 – December 15) and late winter (March). We used a dataset of movements by GPS-collared wolves and cougars to fit the models. We used generalized additive models (GAMs) to estimate the (squared) displacement rate as a smooth, circular function of the time of day. We fit separate models for early and late winter because of changes in time of sunrise and sunset between these two periods. Each model included a term for the population-level pattern and a term for the individual-level pattern, the GAM analog to a random slopes model in a generalized linear mixed model, thus controlling for individual variation.

From each fitted model, we predicted predator activity patterns by predicting displacement rate at the population level. We made separate predictions for wolves and cougars for early and late winter. The model predicted (squared) displacement rate, and absolute distances varied between wolves and cougars. To make these metrics comparable between the two species, we scaled these predictions linearly to range from -1 to 1 for each predator and in each season. We used these scaled activity patterns to estimate the effect of risk on hourly elk behavioral responses, i.e., the temporal component of the dynamic landscape of risk (dLOR).

### Methods

#### Data Collection

Wolves and cougars were fitted with GPS collars as part of long-term monitoring and research programs by the National Park Service. We used data from wolves for winters ranging from 2001 – 2022, and we used data from cougars for winters ranging from 2001 – 2022. We split the data into early winter (November 15 – December 15) and late winter (March 1 – March 31) because of changes in time of sunrise and sunset between these two periods. We refer to early and late winter as “seasons”. We refer to winters with the calendar year of the late winter period, consistent with other references to “winter” or “year” in this paper.

Median sampling rates for wolves ranged from 1 – 6 h (mean = 2.6 h); 130 winter-seasons had 1-h sampling rates (the most common sampling rate). After data preparation (see below), we used data from 62 wolves and 211 wolf-winter-seasons to fit the models.

Median sampling rates for cougars ranged from 1 – 6 h (mean = 1.8 h); 68 winter-seasons had 1-h sampling rates (the most common sampling rate). After data preparation (see below), we used data from 25 cougars and 117 cougar-winter-seasons to fit the models.

#### Data Preparation

We used the data cleaning functions from the R package ‘amt’ for basic cleaning prior to displacement rate calculations. We first kept only those individuals with at least 150 locations per winter-season. We excluded any GPS locations with low precision, which we defined as GPS locations with dilution of precision (DOP) > 6. We removed locations that represented unreasonable speeds with the function `flag_fast_steps()` with argument `delta = 50,000` m^2^/s for both species.

Because different individuals had different sampling rates, we used squared displacement rate (SDR) as the metric to quantify activity. Briefly, expected displacement only scales linearly with time under ballistic (straight line) motion. Under a pure random walk, the square of expected displacement scales linearly with time, thus squared displacement rate, which we measured in m^2^/s, would be expected to be sampling-rate invariant under a random walk. By using SDR, we assumed that predator motion was more similar to a random walk than to ballistic motion.

#### Modeling

We fitted a Generalized Additive Model (GAM) for each predator to estimate predator SDR as a function of time of day. GAMs use penalized splines to represent smooth predictors, thus the relationship between a predictor and the response variable can take a non-linear form that is not pre-specified by the user, and cubic regression splines (and their generalizations) used in GAMs can theoretically represent any smooth function (Wood 2017). This is useful in this context because daily activity patterns are expected to be cyclical and smoothly varying; an arbitrary smooth function can capture this without specifying daily breaks a priori (e.g., sunrise, sunset, dusk, etc.). The penalty (as in “penalized splines”) reduces the total degrees of freedom used by each smooth term; i.e., the data inform how much complexity a spline should have, and as long as the user provides enough basis functions a priori, the model will select an appropriate number of basis functions to parsimoniously represent the relationship. The penalty induces shrinkage exactly analogous to the shrinkage of random effects in a generalized linear mixed model (Wood 2017). For that reason, GAMs can also be used to flexibly fit random effects.

The four models (two predators, two seasons) had the same form. The response variable was the natural logarithm of SDR (hereafter, log-SDR), modeled as a normal random variable. The predictor variable was hour of the day, which we assigned as the midpoint between consecutive GPS locations. We expressed the relationship between log-SDR and hour using two splines: a circular cubic regression spline that represented the population-level activity pattern and a factor spline that allowed the pattern to vary by individual. Pedersen et al. (2019) refer to this type of formulation as “Model GS”, and they explain in detail why this model is analogous to a generalized linear mixed model with random slopes for each individual. The circular cubic regression splines ensured that predicted values at the beginning and ending hours of each day matched; i.e., since hour 0 and hour 24 both represent midnight, the model prediction should match.

Previous research in this system, with a smaller sample size, found evidence of different activity patterns for male and female cougars (Kohl *et al*. 2019). All cougar data included in this previous model were included here (6 individuals), but our sample size here was much larger (25 individuals). We explored a model for cougars where the population-level pattern included an effect of sex (via a tensor product interaction) and a sex-specific intercept. While the intercept for females was lower than that of males, the relative activity patterns showed no evidence of being different from the population-level activity pattern (ANOVA; females, p = 0.56; males, p = 0.71). We presented the model predictions from this model below, which show the shape of the patterns are the same between sexes, although females move shorter distances.

#### Predicting Predation Risk

We used the four fitted models to predict displacement rate in km/h. We used only the population-level splines to make the prediction, thus controlling for, but not predicting with, individual variation. The raw model predictions were in log-SDR, which we exponentiated to convert to SDR. We used the SDR to calculate the expected displacement in one hour using the `amt` function `get_displacement()`, which returned the expected displacement in m. We converted to km, and interpreted the result as km/h.

Wolves and cougars had different movement patterns, and so the absolute magnitudes of displacement rate were different between the two predators. The goal of this was not to estimate absolute displacements, but rather relative activity patterns. To facilitate comparison between the two species and across seasons, we linearly scaled the displacement rates to range from -1 to 1.

### Results

In both seasons, wolves had higher peaks in expected displacement than did cougars (Fig. S2.1), consistent with the more cursorial behavior of wolves.

**Figure S2.1.**
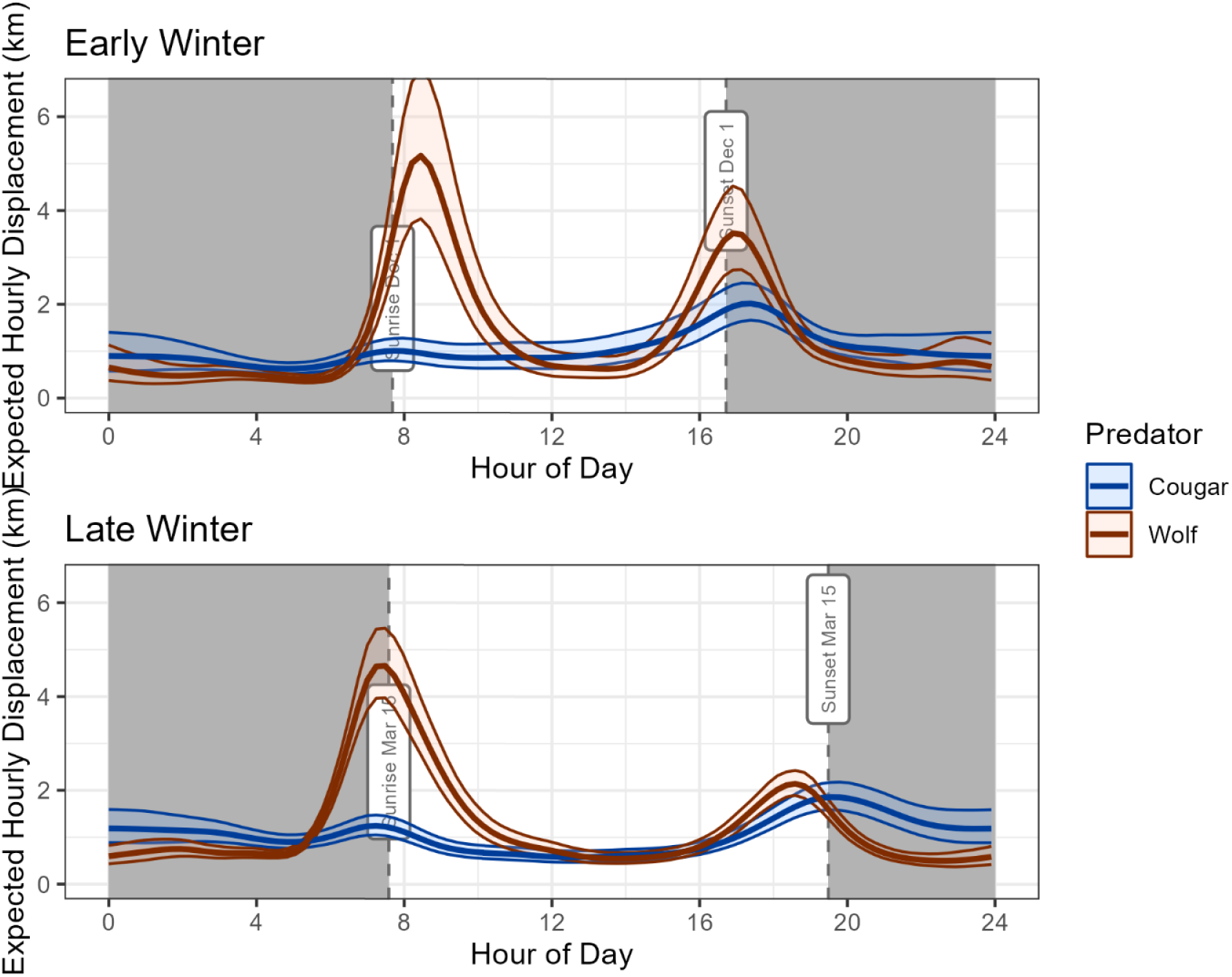
Predator displacement rates. We used the fitted models to predict predator displacement rates as the expected displacement (in km), given a one-hour time step. Wolves had higher displacement rates than cougars, somewhat obscuring the relative patterns. Top panel shows early winter (November 15 – December 15) and bottom panel shows late winter (March 1 – March 31). Envelopes show 95% confidence interval. Shaded boxes show nighttime, with dashed vertical lines showing sunrise and sunset times in Gardiner, MT, for the median date of that season (December 1 or March 15).

The activity index showed the relative patterns between wolves and cougars more clearly (Fig. S2.2). In both seasons, wolves had their greatest activity near sunrise, with a secondary peak near sunset. Cougars showed the opposite pattern, with the greatest activity near sunset and a secondary peak near sunrise. Cougars showed more nocturnal activity than wolves in both seasons. We used the values of activity index shown here (Fig. S2.2) to quantify temporal variation in predation risk.

**Figure S2.2.**
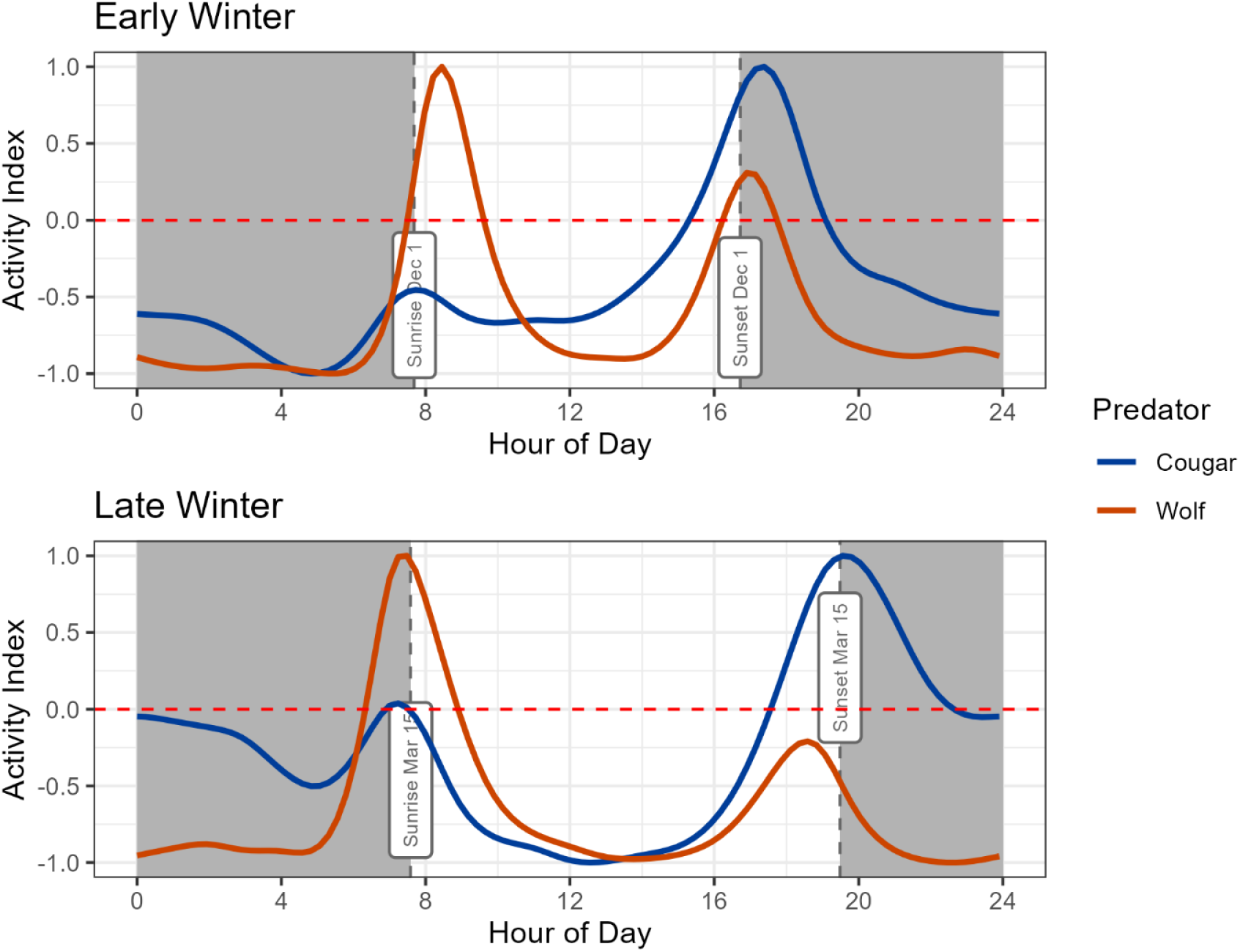
Predator activity index. We linearly scaled the predicted hourly displacement to range from -1 to 1 for each predator in each season, better showing relative activity patterns. Wolves were most active near sunrise, with a secondary peak near sunset; cougars showed the opposite pattern, as well as more nocturnal activity. Shaded boxes show nighttime, with dashed vertical lines showing sunrise and sunset times in Gardiner, MT, for the median date of that season (December 1 or March 15). Values shown here were used to quantify temporal variation in predation risk.

Cougars showed the same activity pattern throughout the day in both seasons, regardless of sex (Fig. S2.3). Because we showed the effect of sex was not important, we used the simpler model presented above to quantify temporal variation in cougar risk.

**Figure S2.3.**
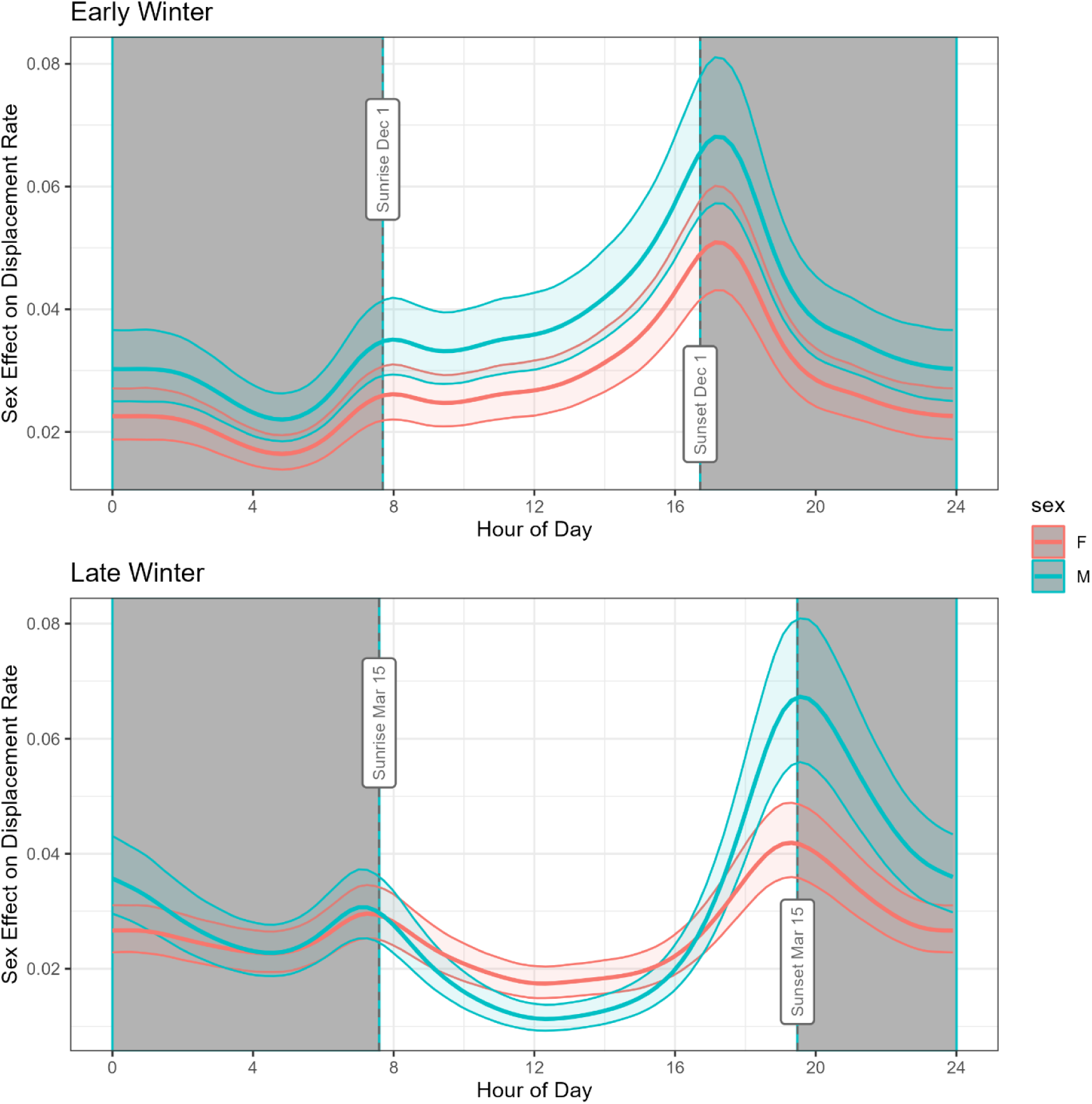
Cougar sex effect. Because previous research suggested male and female cougars may have different activity patterns, we fitted a cougar model to test for this. There was no significant difference in activity patterns between male and female cougars, although female cougars moved less in an absolute sense (captured by a sex-specific intercept). Envelopes show 95% confidence intervals. Shaded boxes show nighttime, with dashed vertical lines showing sunrise and sunset times in Gardiner, MT, for the median date of that season (December 1 or March 15).

## Appendix S3. Additional iSSA description

The first-stage model was an integrated step selection analysis (iSSA), with individual integrated step-selection functions (iSSFs) fitted to each elk-winter-season of data. Covariates in the iSSA are described synthetically in the main text (Table 1), but here, we expand on the detail.

### Model Formula

We modeled habitat selection as a function of resources, risks, and conditions (Matthiopoulos et al. 2015). We allowed movement to vary as a function of snow-water equivalent and time of day, and we allowed for correlation between step lengths and turn angles. The full model specification, as an R model formula, was:

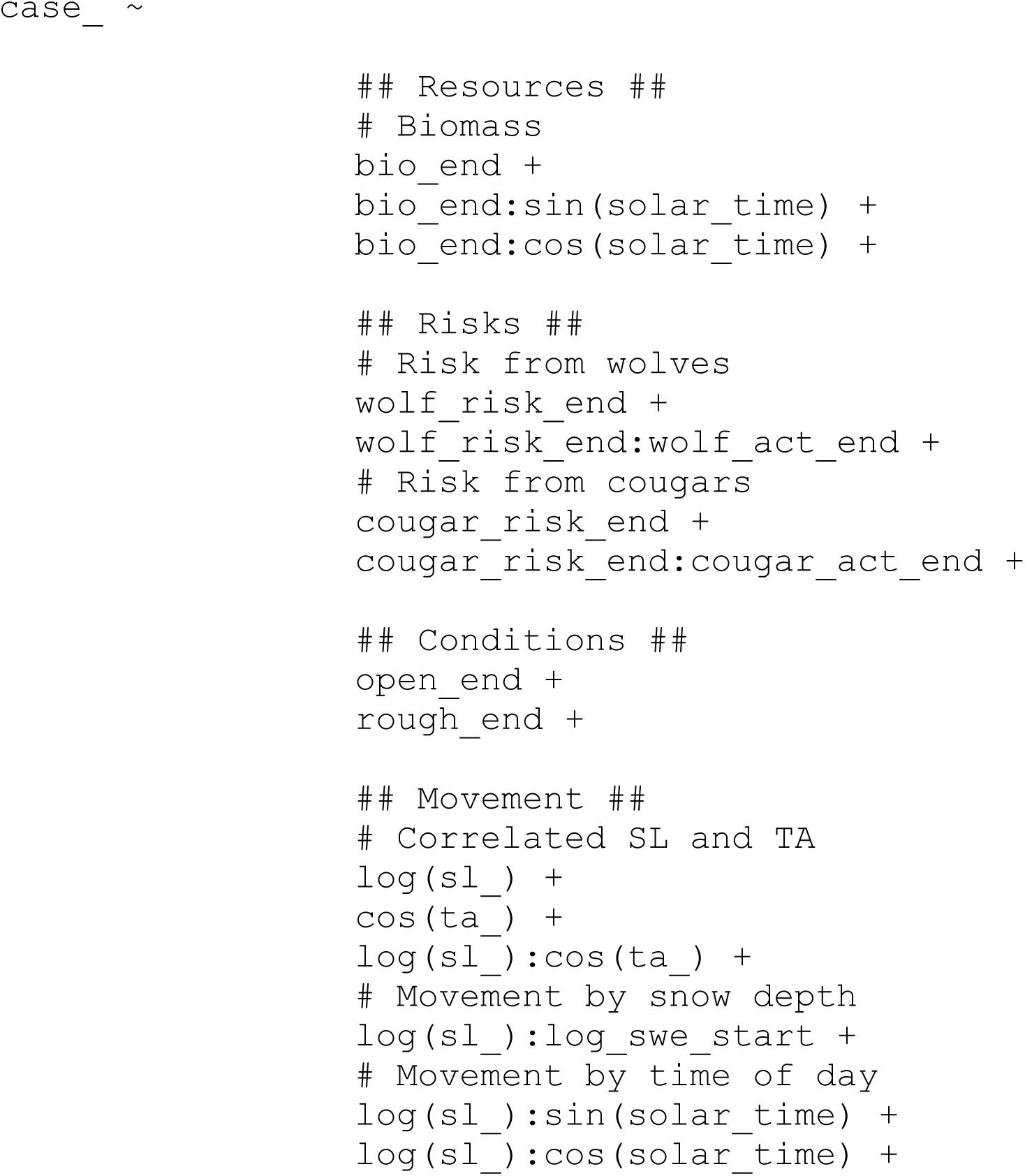
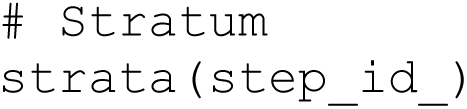

Variable names with `_end` appended indicate the covariate was attached to the endpoint of the step, whereas names with `_start` appended indicate the covariate was attached to the start point of the step. The variable `case_` is the response variable, indicating whether a step was used (1) or available (0).

The model is fit via conditional logistic regression, which is stratified such that each used step is compared only to its corresponding available steps with the same start point. The variable `step_id_` pairs together those corresponding used and available steps, and the special function `strata()` indicates that to the model fitting routine.

#### Resources

We modeled resources using herbaceous biomass (`bio`), using the herbaceous biomass product from the Rangeland Analysis Platform (Jones et al. 2021; Robinson et al. 2019; Smith et al. 2023). We included an interaction between biomass and sine and cosine transformations of solar time of day, which allowed elk foraging behavior to vary across the diel cycle and with changes to daylength. We describe solar time below.

#### Risks

We modeled risks using the dynamic landscape of risk (dLOR), which we described in detail in Appendix S1 and S2. The variable names with `_risk_` describe risky places and the variable names with `_act_` describe risky times (predator activity). The interaction between risky places and risky times creates the dLOR.

#### Conditions

We modeled conditions with canopy openness (`open`) and terrain roughness (`rough`), which have been used to model risky places in this system, and they are included our risk layers (Appendix S1). We included these landscape variables in the iSSA because elk may also select them for other reasons (see main text).

#### Movement

We modeled movement using a Gamma distribution for step lengths and a von Mises distribution for turn angles. The iSSA works by sampling available steps from parametric “tentative” distributions and then updating the parameters of those distributions using coefficients from the model (Avgar et al. 2016). In the case of the Gamma distribution, the coefficient for log of step length (`log(sl_)`) updates the shape parameter and the coefficient for step length (`sl_`) updates the scale parameter. In the case of the von Mises distribution, the coefficient for cosine of turn angle (`cos(ta_)`) updates the concentration parameter. Interactions with other variables express these parameters of the movement distribution as functions of the interacting variable (reviewed in Fieberg et al. 2021). Thus, we allowed the step length distribution to vary with the log of snow-water equivalent (`log_swe`) and with the solar time of day (`solar_time`; described below). We estimated snow-water equivalent in each pixel with the SWE product from Daymet 4.0 (Thornton et al. 2022).

#### Solar Time of Day

While we modeled diel pattern in the dLOR using predator activity (Appendix S2), for all other variables, we modeled diel pattern using sine and cosine transformations of the solar time of day. We used the function `getSunlightTimes()` from the R package `suncalc` to retrieve the precise times of solar passage for a specific location (the start point of the step) at a specific time (the start point of the step). We calculated the time of the solar nadir (sun directly under foot), sunrise, solar noon, and sunset. For each day, we evenly spaced values from −*π* to *π* to correspond to these solar positions (`solar_time`): nadir to sunrise spanned −*π* to 0, sunrise to noon spanned 0 to *π*, noon to sunset spanned *π* to 0, sunset to nadir spanned 0 to −*π* (Fig. S3.1A). We took the cosine (`cos(solar_time)`; Fig. S3.1B) and sine (`sin(solar_time)`; Fig. S3.1C) of these values to use in interactions in our model. These transformations of solar time interacted with biomass and the step length distribution.

**Figure S3.1.**
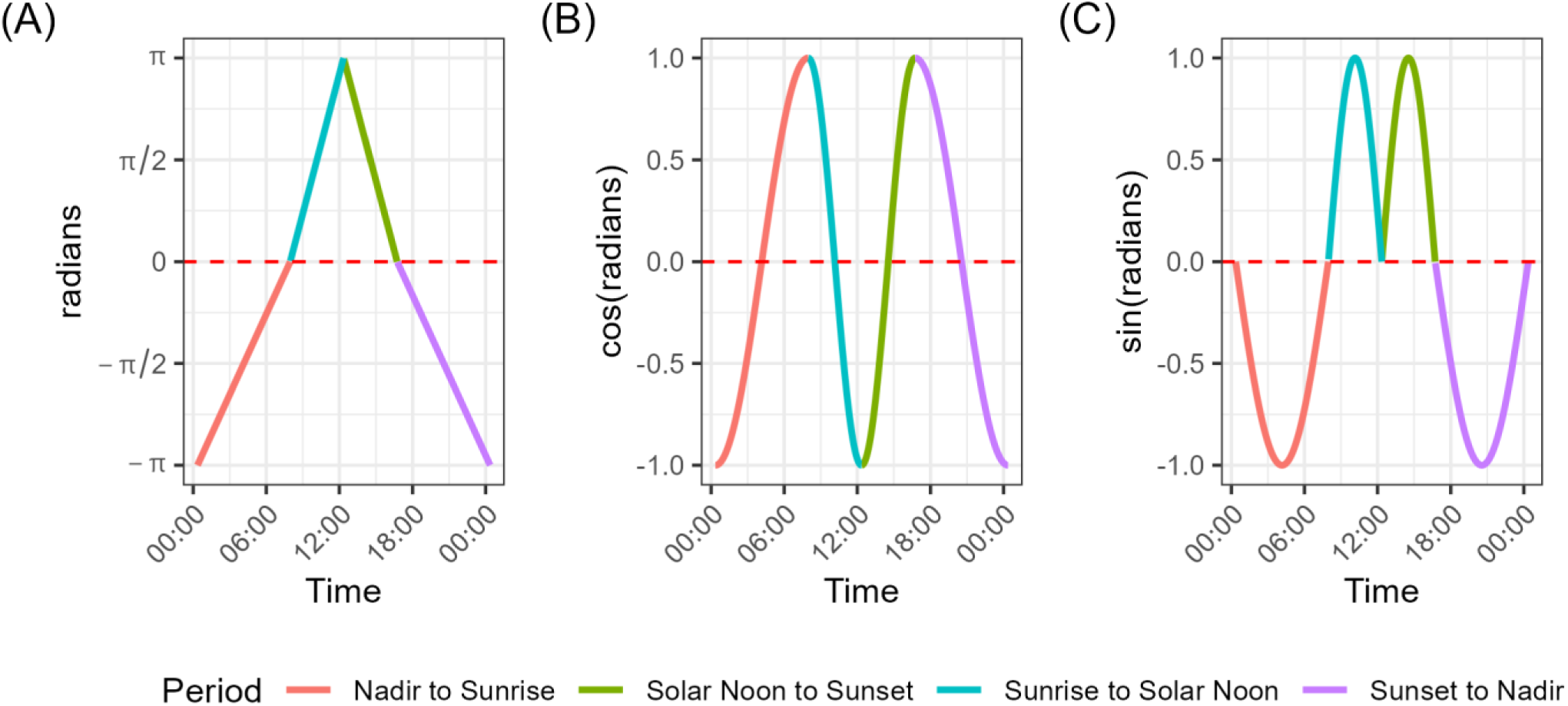
Solar time of day. For a day and location (the start of the step), we calculated the precise timing of the solar nadir, sunrise, solar noon, and sunset. We evenly spaced values from −*π* to *π* to correspond to these solar positions: nadir to sunrise spanned −*π* to 0, sunrise to noon spanned 0 to *π*, noon to sunset spanned *π* to 0, sunset to nadir spanned 0 to −*π* (A). We took the cosine (B) and sine (C) of these solar times to use in interaction with biomass and the step-length distribution. These example plots show times for 21 December 2020 at 44.98101°N, 110.4089°W.

### Estimating Risk Taking (Defensive Trait)

We used the fitted iSSFs to estimate the defensive trait for each elk-year-season using relative selection strengths (Avgar et al. 2017). We did this by making a hypothetical comparison between a high-risk/high-reward habitat (HRH) and a low-risk/low-reward habitat (LRH). In this case, the RSS gives how many times more likely an elk is to select the HRH over the LRH, which is meant to provide a relatively intuitive interpretation of this complex model. We defined HRH as a habitat with the 90th quantile of biomass and the 90th quantile of risk and LRH as a habitat with the 10th quantile of biomass and the 10th quantile of risk.

Only the magnitude, not the sign or significance, of the effect was sensitive to the quantiles of forage and risk we chose – the sign and significance are determined by the fitted coefficients from the model. To see this, consider the definition of RSS. In general, the log-RSS for two habitats (*x*_1_, *x*_2_) that differ in a single habitat variable (ℎ*_i_*) is given by (Avgar et al. 2017):

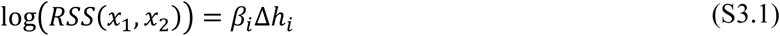

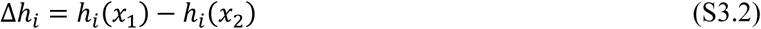

That is, it is the estimated coefficient (*β_i_*) times the difference in the values of the habitat covariate (Δℎ*_i_*) in the two habitats. In our case, the HRH and LRH differ in both resources (herbaceous biomass) and risks (dLOR), so the log-RSS is the sum of both the habitat variable effects, e.g.,

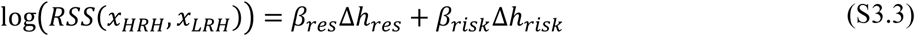

But note that because there are interactions with time-varying covariates for both resources and risks, the actual formula contains more terms.

Eqn. S3.3 shows why the inference is not sensitive to the choice of quantile of resources/risks for the HRH and LRH. The choice of quantile (e.g., 90^th^ and 10^th^ vs. 80^th^ and 20^th^ vs. 60^th^ and 40^th^) scales Δℎ up or down, but it would not change its sign (unless high and low were reversed, e.g., 90^th^ and 10^th^ vs. 10^th^ and 90^th^, which is constrained by definition). Thus, sign and significance are determined by the point estimate and uncertainty in the coefficients, not the choice of the quantiles.

Note that using different quantiles for resources vs. risks would be possible. We chose to use the same quantiles to equally weight the selection of resources and selection (avoidance) of risks.

## Appendix S4. Supplemental Tables

**Table S4.1.** Predicted population-mean risk-taking behavior. We calculated predicted risk-taking behavior for elk aged 1 – 20 using the second stage model (see Fig. S5.1 and main text). We estimated the proportion of elk in each age class (1 – 20) using the reconstruction of Hoy et al. (2019; see main text) for years in which the elk population was at its youngest (1995, median age = 4 y) and at its oldest (2009, median age = 10 y). We then took the mean of the risk-taking behavior, weighted by the proportion of the population in each age class, for the two age structures. We did this across all bootstrap iterations to propagate uncertainty, and we present the mean and 95% confidence intervals across the bootstrap iterations. We present the results on the log scale (log-RSS) and the natural scale (RSS). The column “Percent Change” shows the decrease in risk-taking behavior (RSS) from 1995 (young age structure) to 2009 (old age structure) as a percentage of the 1995 behavior. We did not present percent change for cougars because there was no significant effect of age, and the confidence intervals include extreme, nonsensical values.

| Predator | Time | Year | log-RSS (95% CI) |  |  | RSS (95% CI) |  |  | Percent Change (95% CI) |  |  |
| --- | --- | --- | --- | --- | --- | --- | --- | --- | --- | --- | --- |
|  |  |  | Mean | Lower | Upper | Mean | Lower | Upper | Mean | Lower | Upper |
| Cougar | Riskiest | 1995 | -2.64 | -8.58 | 4.89 | 204.90 | 0.00 | 133.43 | N/A |  |  |
|  |  | 2009 | -2.69 | -7.42 | 4.65 | 116.42 | 0.00 | 104.35 |  |  |  |
|  | Mean | 1995 | 1.42 | -2.07 | 7.23 | 1868.01 | 0.13 | 1383.38 | N/A |  |  |
|  |  | 2009 | 1.73 | -0.90 | 4.00 | 12.79 | 0.41 | 54.35 |  |  |  |
|  | Safest | 1995 | 2.61 | -3.99 | 7.84 | 5963.80 | 0.02 | 2542.42 | N/A |  |  |
|  |  | 2009 | 2.59 | -3.99 | 6.84 | 138.82 | 0.02 | 931.11 |  |  |  |
| Wolf | Riskiest | 1995 | 0.93 | 0.10 | 1.80 | 2.79 | 1.10 | 6.04 | 41% | 23% | 58% |
|  |  | 2009 | 0.38 | -0.28 | 1.06 | 1.56 | 0.76 | 2.90 |  |  |  |
|  | Mean | 1995 | 1.38 | 0.80 | 1.97 | 4.14 | 2.24 | 7.15 | 42% | 28% | 52% |
|  |  | 2009 | 0.82 | 0.38 | 1.27 | 2.34 | 1.46 | 3.57 |  |  |  |
|  | Safest | 1995 | 1.61 | 0.93 | 2.28 | 5.32 | 2.53 | 9.82 | 42% | 25% | 55% |
|  |  | 2009 | 1.05 | 0.55 | 1.59 | 2.95 | 1.73 | 4.89 |  |  |  |

## Appendix S5. Supplemental Figures

**Figure S5.1.**
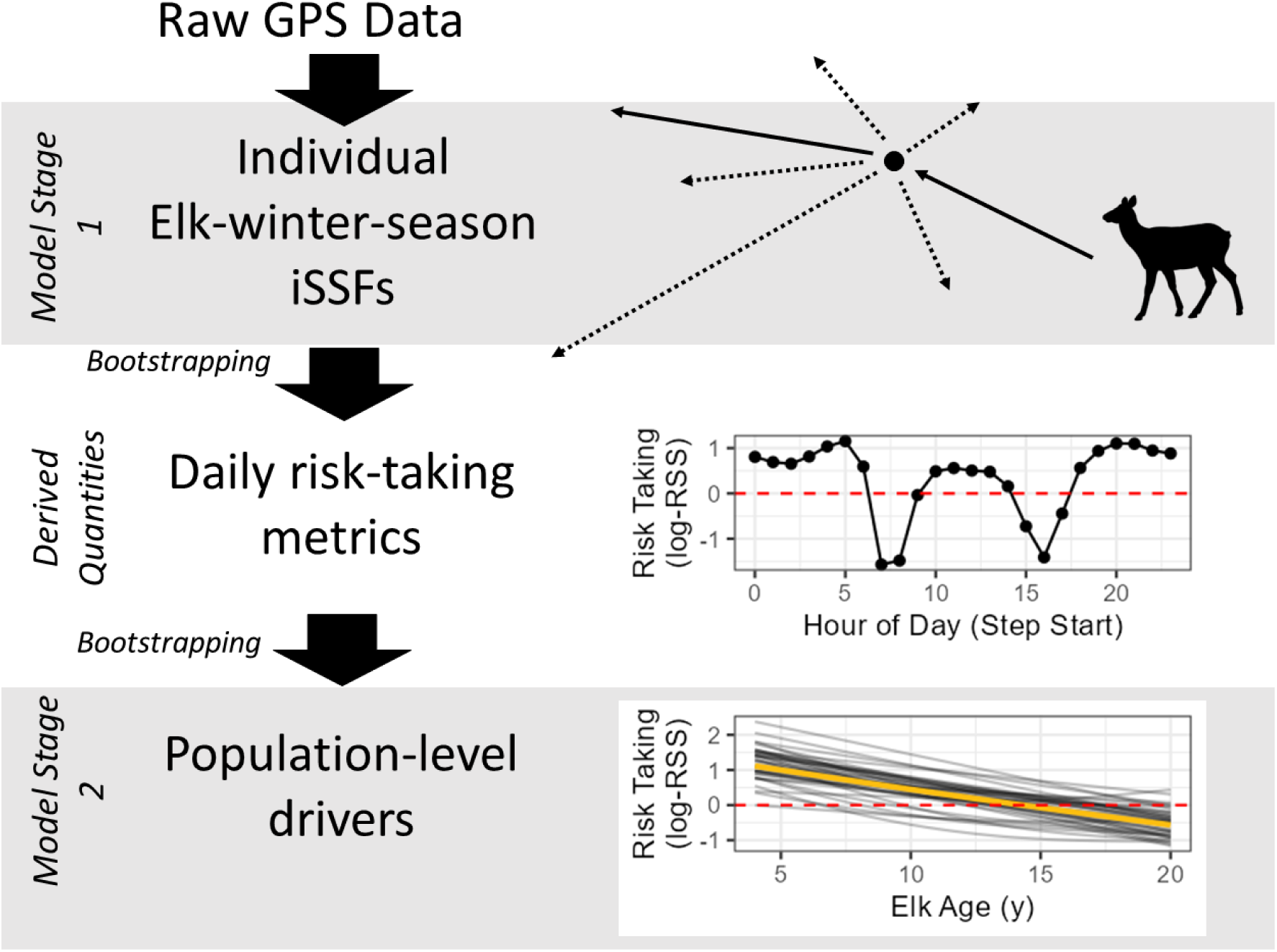
Conceptual figure showing the workflow starting with raw GPS data, the first-stage model (iSSA), the derived risk-taking behavior, and the second-stage model (population-level inference). The first-stage model was an integrated step selection analysis (iSSA), with individual integrated step-selection functions (iSSFs) fitted to each elk-winter-season of data (see main text and Appendix S3 for predictor variables). The daily risk-taking metrics (derived quantities) were calculated for each elk-winter-season from the corresponding fitted iSSF, with uncertainty propagated through parametric bootstrapping. The daily risk-taking metrics were calculated as the log-relative selection strength (log-RSS) for a hypothetical choice between a high-risk/high-reward habitat and a low-risk/low-reward habitat, with positive values indicating a preference for the high-risk/high-reward habitat and negative values indicating a preference for the low-risk/low-reward habitat. The second-stage model quantified the population-level pattern in risk taking while controlling for alternative drivers (see main text). We formulated these models using generalized additive models (GAMs) to allow for potential non-monotonic patterns. We fitted separate second-stage models for each combination of predator (wolf or cougar) and time of day (maximum predator activity, minimum predator activity, daily mean). We fitted models for each bootstrap iteration returned from stage 1 to propagate the uncertainty in the iSSA through to the population-level models.

**Figure S5.2.**
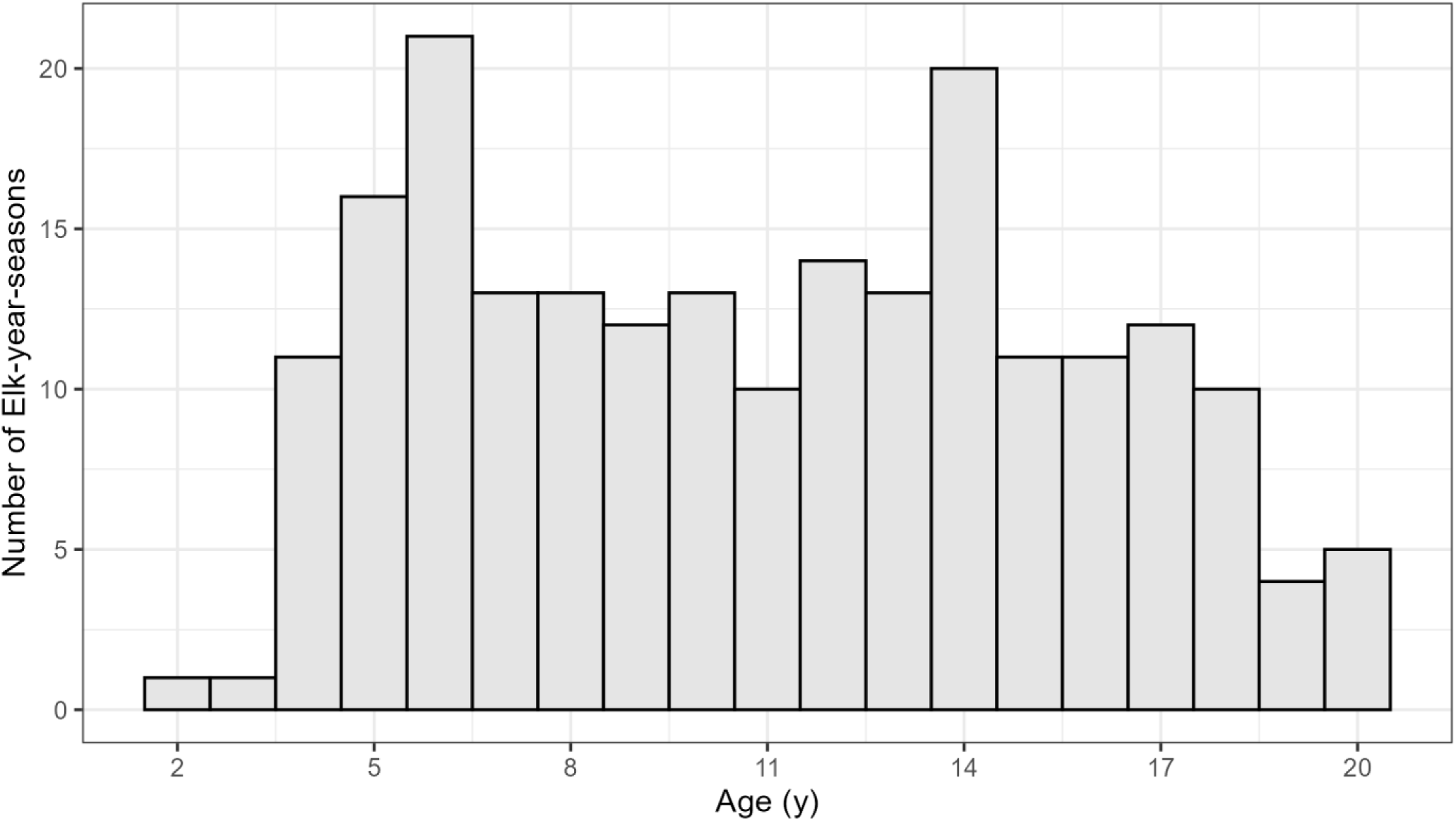
Histogram of ages for each elk-year-season. Our fitted models included 214 elk-year-seasons from 72 individuals across 5 years (2016 – 2020). Ages ranged from 1.5 to 20.5 y.

**Figure S5.3.**
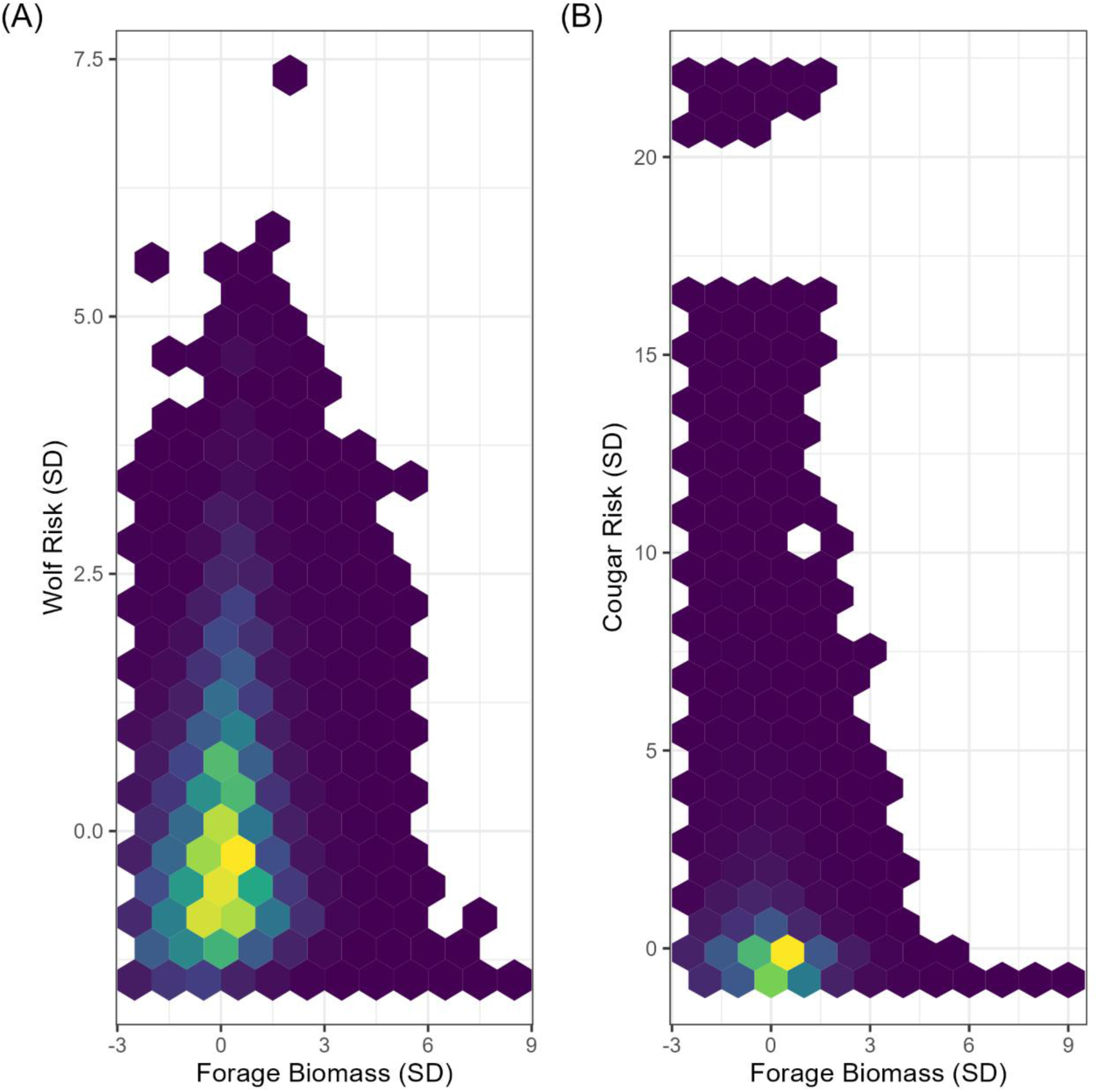
Correlation between forage and risk. Hexbin plot shows the number of data points that fall within each bin, with dark colors representing few points, bright colors representing many points, and empty cells indicating no data points. We calculated correlation between forage (x-axis) and risk (y-axis) using the available steps from the integrated step selection analysis, thus defining the correlation for only the domain of availability for this analysis. Correlation between forage biomass and wolf risk was weakly positive (Spearman’s r = 0.17; A). Correlation between forage biomass and cougar risk was weakly negative (Spearman’s r = -0.16; B). Units of both forage and risk are standard deviations (SDs) because we scaled and centered the covariates with a z-transformation.

**Figure S5.4.**
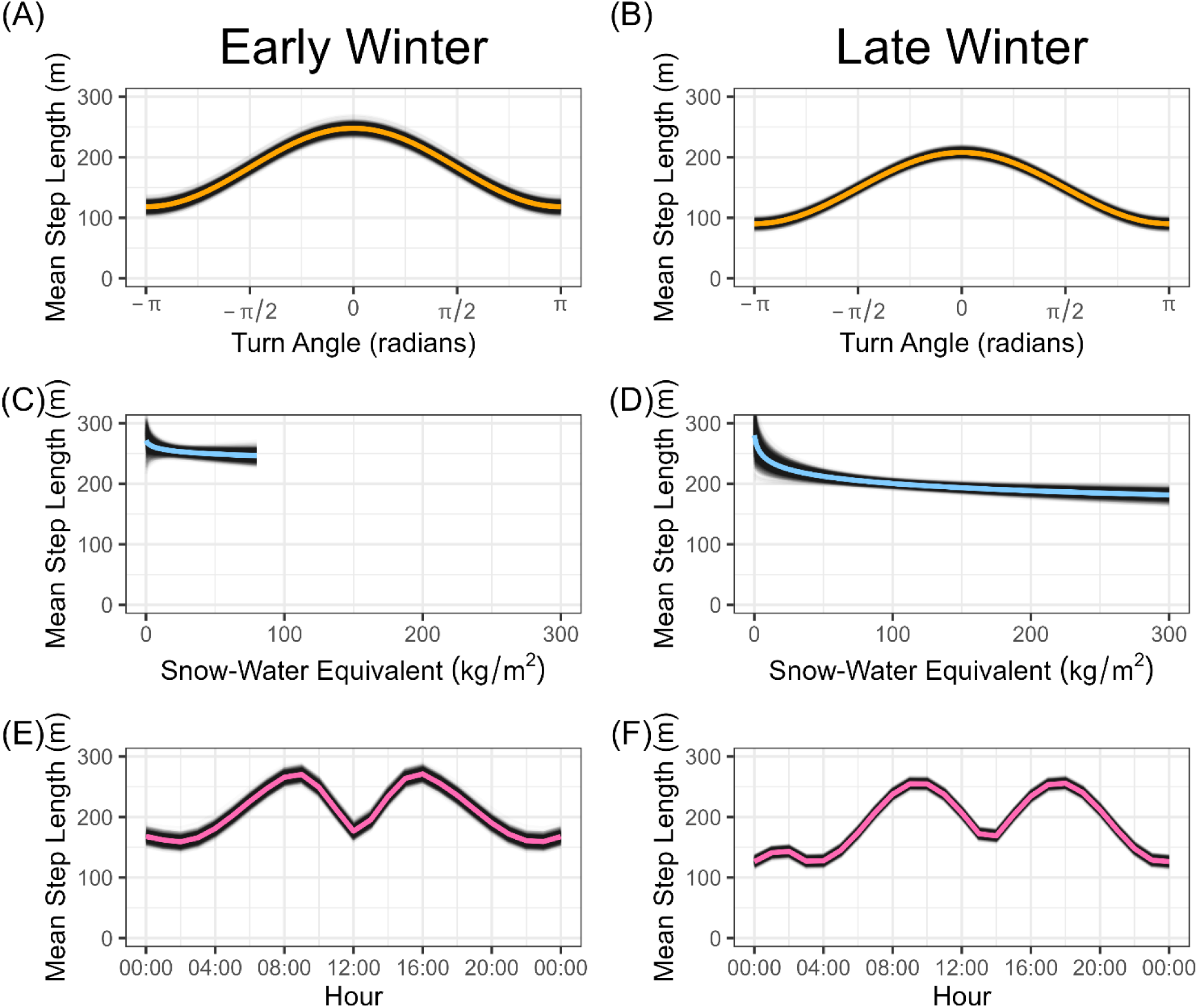
Effect of covariates on movement model. Our integrated step selection functions (iSSFs) included interactions in the movement part of the model allowing for a dynamic step length distribution; here, we show the mean step length from the fitted Gamma distribution. Mean step lengths were larger in early winter (A) and late winter (B) when turn angles were close to 0. Snow-water equivalent (SWE) decreased mean step lengths in both early winter (C) and late winter (D); note that maximum observed SWE was much greater in late winter than early winter and the curve in (C) is drawn accordingly. Mean step length also varied with time of day, with peaks in the morning and evening in both early winter (E) and late winter (F). For all panels, all other covariates were held at their mean. Early winter is represented by December 1 and late winter is represented by March 15. Thick lines show the mean across all bootstrap iterations (n = 2000), while the thin, transparent lines show bootstrap iterations.

**Figure S5.5.**
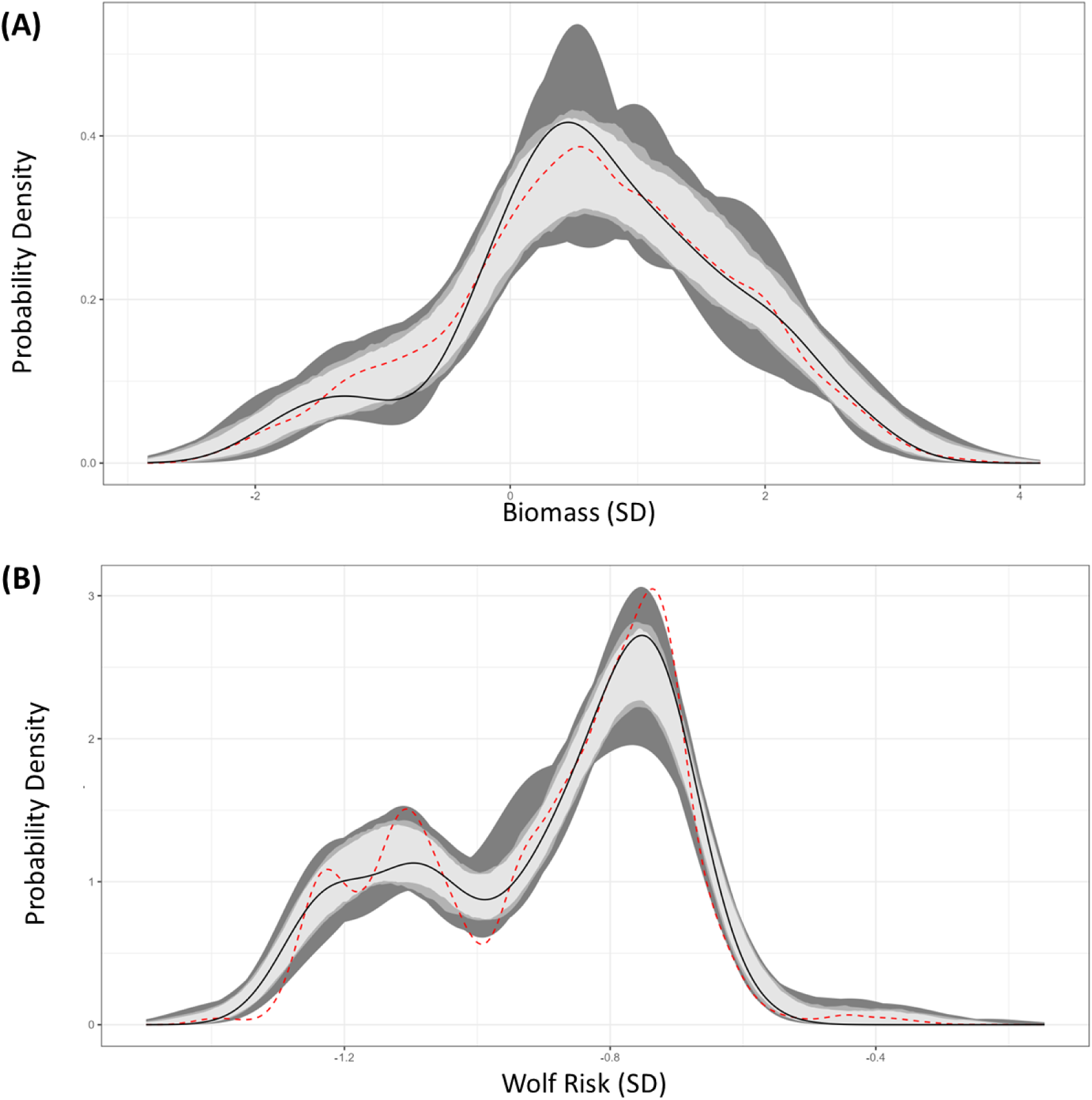
Example used-habitat calibration plots (UHC plots). We created UHC plots by using parametric bootstrap resampling. UHC plots show the distribution of each habitat variable for the used steps (black line), available steps (red, dashed line), and fitted model steps (gray envelopes). We created model distributions by using parametric bootstrapping to account for uncertainty. For each iteration of the bootstrap, we resampled the parameters of each integrated step-selection function from a multivariate normal distribution. We used the bootstrap samples to create 90%, 95%, and 100% empirical confidence envelopes (gray shading). When the distribution of habitat for modeled steps agrees with the distribution of habitat for used steps, the model is well-calibrated (black line falls inside the gray envelope). The difference between the available (red, dashed line) and used (black line) distributions depicts habitat selection. Shown here are two example UHC plots for one individual elk-winter-season. The top panel shows a UHC plot for herbaceous biomass (A). The bottom panel shows a UHC plot for wolf risk, accounting for dynamic habitat selection (B). UHC plots for all habitat variables across all individual elk-winter-seasons showed high calibration.

**Figure S5.6.**
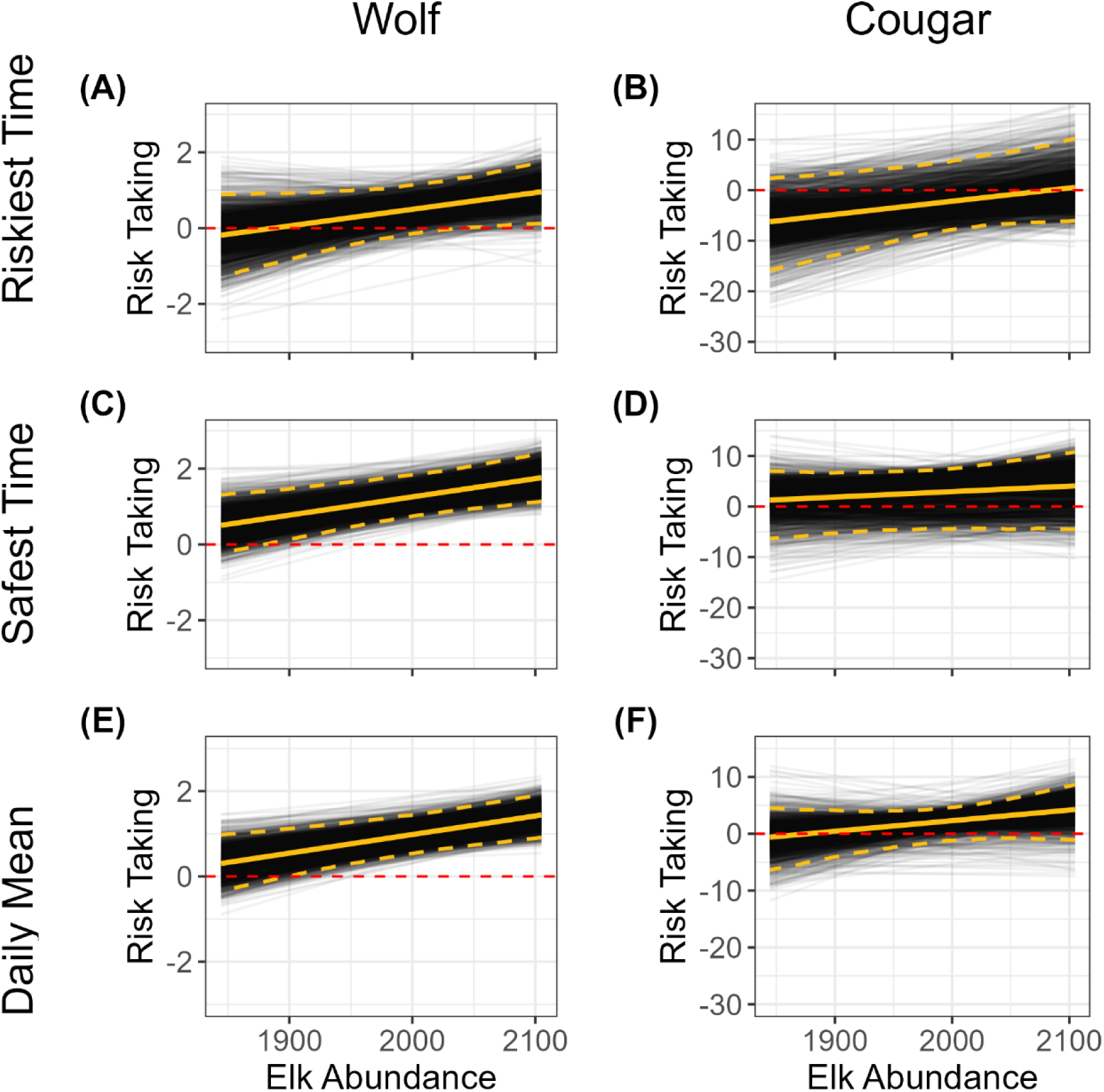
The population-level effect of elk density on risk taking, estimated from bootstrap samples (n = 2000). Elk risk taking increased slightly with elk density with respect to both wolves (left column) and cougars (right column), as expected due to various safety in numbers effects.

**Figure S5.7.**
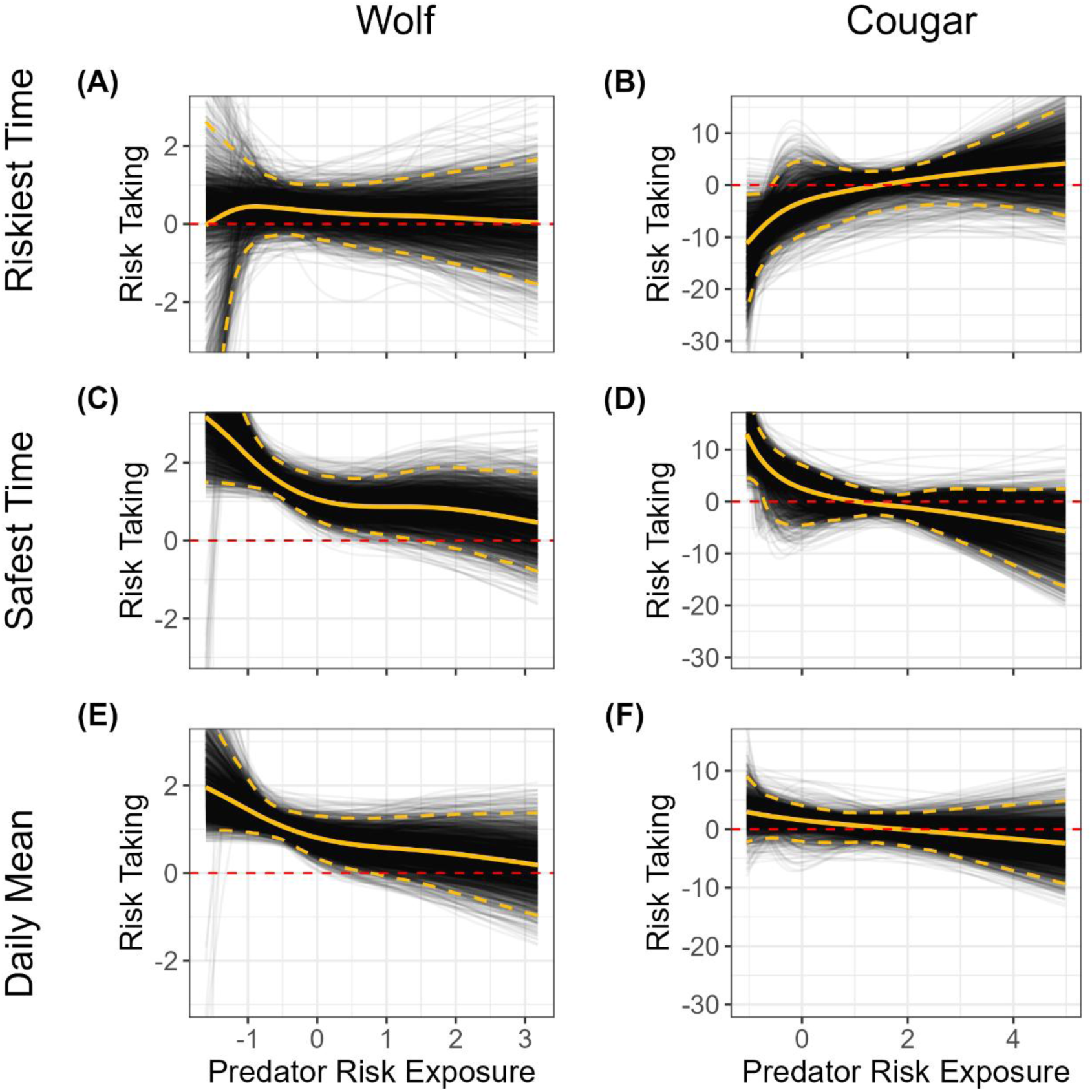
The population-level effect of predator risk exposure on risk taking, estimated from bootstrap samples (n = 2000). We found highly uncertain effects of risk exposure on risk taking during the riskiest time (A – B). With respect to both predators, safest time risk taking decreased with risk exposure (C – D), i.e., elk exposed to more predation risk played it safer, even at the safest time of day. On average throughout the day, increased exposure to wolf risk resulted in decreased risk taking (E), whereas increased exposure to cougar risk had little effect (F).

**Figure S5.8.**
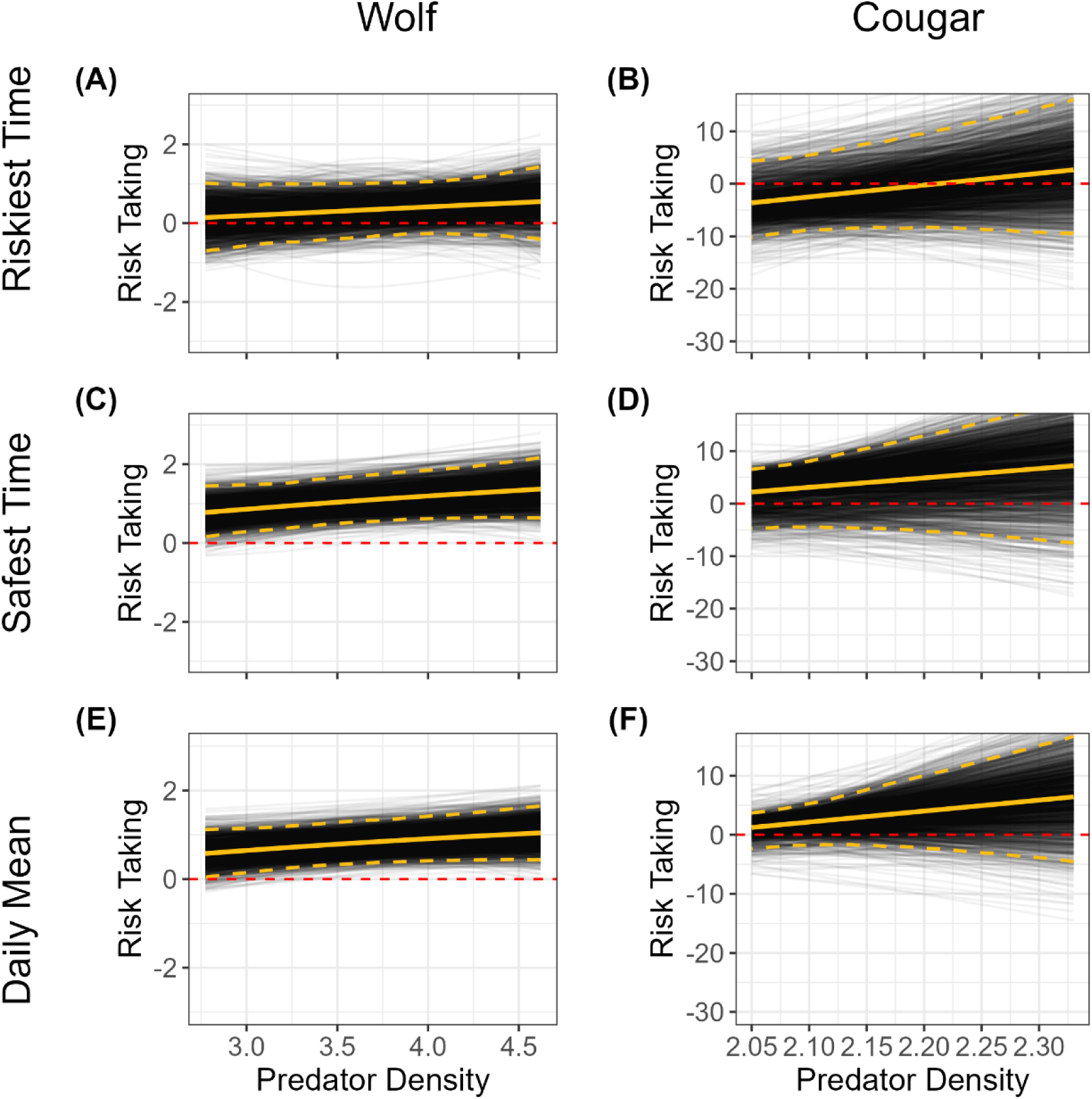
The population-level effect of predator density on risk taking, estimated from bootstrap samples (n = 2000). There was no consistent relationship between elk risk taking and wolf density (left column) or cougar density (right column).

**Figure S5.9.**
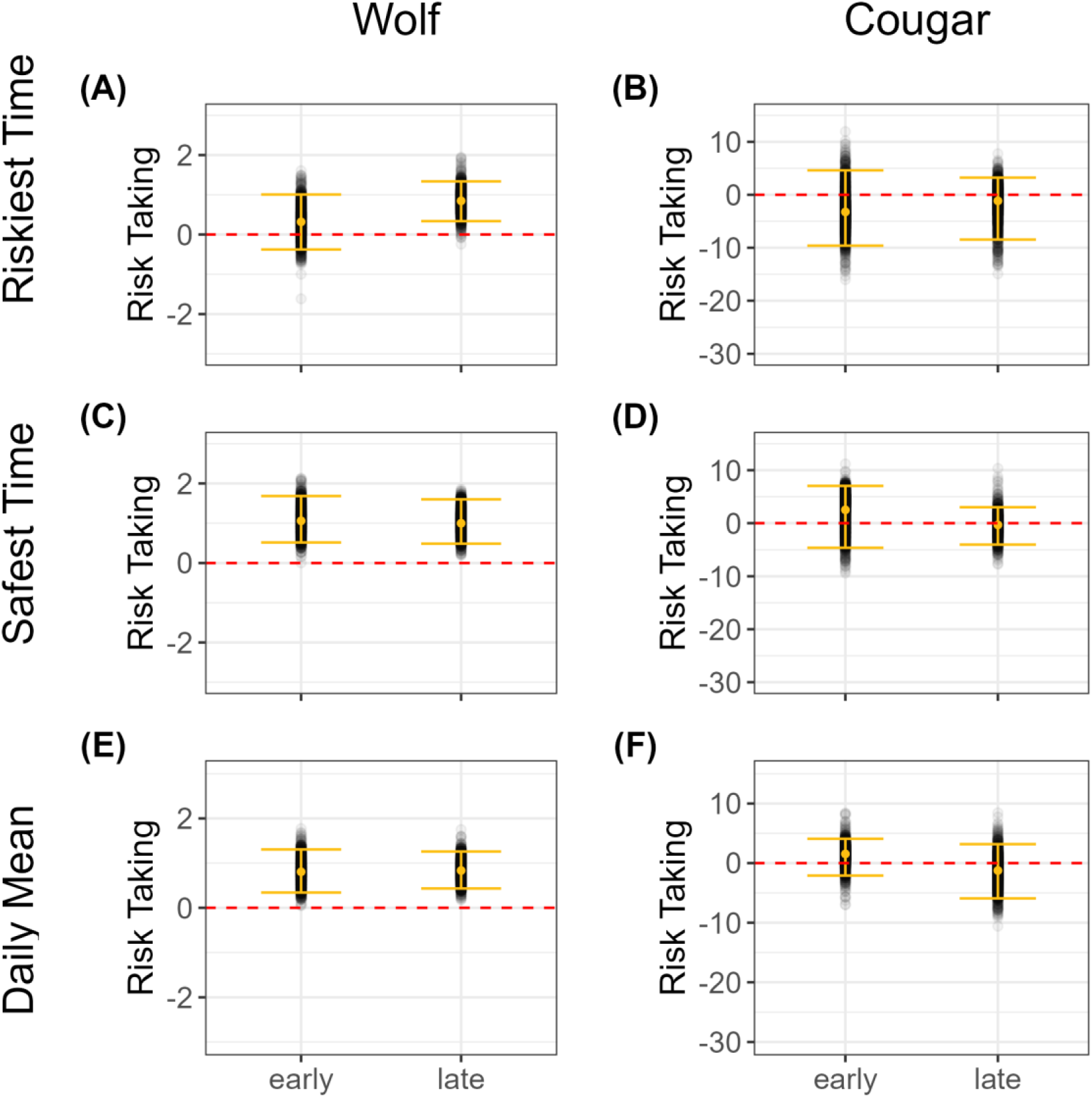
The population-level effect of season on risk taking, estimated from bootstrap samples (n = 2000). There was no consistent relationship between elk risk taking and season with respect to wolves (left column) or cougars (right column).

**Figure S5.10.**
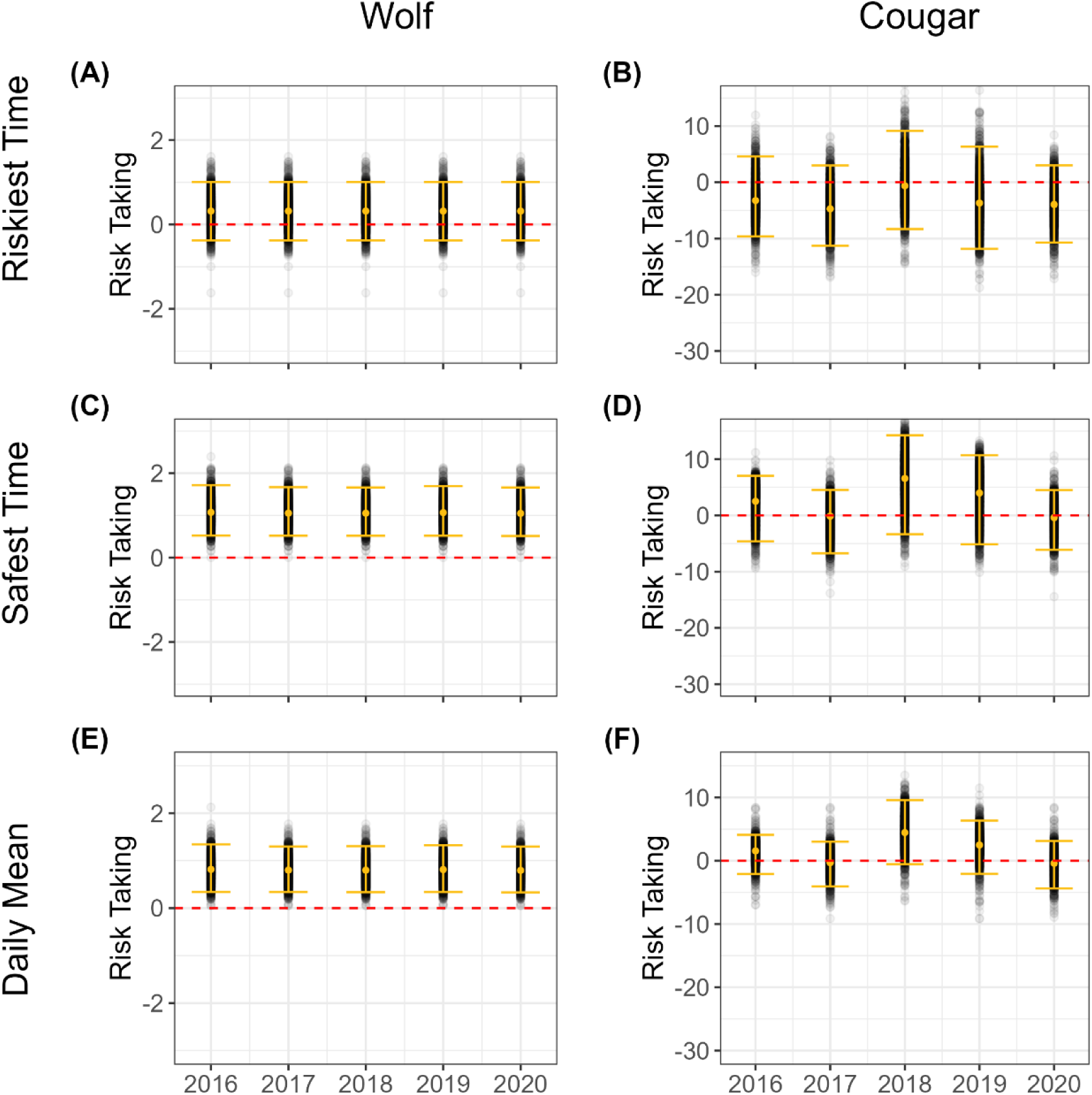
The population-level effect of year on risk taking, estimated from bootstrap samples (n = 2000). There was no clear difference in elk risk taking across years with respect to wolves (left column) or cougars (right column).

